# Multidimensional Cellular Micro-Compartments to Model Invasive Lobular Carcinoma Dormancy

**DOI:** 10.64898/2026.01.09.698501

**Authors:** Xilal Y. Rima, Sarmila Majumder, Chunyu Hu, Divya S. Patel, Hong Li, Xin Huang, Kim Truc Nguyen, Jacob Doon-Ralls, Chiranth K. Nagaraj, Mangesh D. Hade, Setty M. Magaña, Eswar Shankar, Xiaoli Zhang, Daniel G. Stover, Bhuvaneswari Ramaswamy, Eduardo Reátegui

## Abstract

Invasive lobular carcinoma (ILC) accounts for 10-15% of breast cancers. Despite favorable responses to anti-estrogen therapy, the dissemination of cancer cells and resistance to therapies are significant risks for patients with ILC. Late recurrences are prevalent in ILC, suggesting that disseminated tumor cell dormancy may be a mechanism preceding their late overt growth into metastatic lesions. Herein, we investigated the relationship between anti-estrogen resistance and dormancy through multidimensional *in vitro* models. The bioengineered platforms recapitulated the morphological characteristics of ILC and highlighted its distinction from invasive ductal carcinoma. Inducing a dormant phenotype revealed epigenetic changes and enhanced chemical and mechanical sensing of anti-estrogen-resistant ILC cells to the substrate surface, with p27^Kip1^ signaling playing a central role. We propose this platform as a high-throughput method to investigate the propensity of dormancy and its manifestation via a simplified and expedited approach.

## Introduction

The 5-year survival rate of localized breast cancers is >95%,^1^ attributed to advancements in loco-regional therapies, such as surgery and radiation,^2^ as well as effective systemic adjuvant therapies that target key molecular pathways.^3,4^ However, as breast cancer spreads to secondary sites, the 5-year survival rate reduces to ∼32%,^1^ indicating a critical role for disseminated tumor cells (DTCs).^5^ Recurrences can occur after long-term remission, serving as clinical evidence that DTCs acquire a dormant phenotype^6^ and evade conventional therapeutic agents.^7,8^ Late recurrences are prevalent in hormone receptor (HR)-positive breast cancers.^9^ Invasive ductal carcinoma (IDC) and invasive lobular carcinoma (ILC) are the two most common histologic subtypes of HR-positive breast cancer.^10^ Despite comprising 10-15% of breast cancers, ILC is significantly understudied and clinically treated based on evidence from IDC research^11^ notwithstanding its distinct morphological,^12,13^ biomolecular,^14,15^ and recurrence patterns.^16-18^ Patients with ILC face lower rates of long-term disease-free and overall survival,^19,20^ and have an increased number of circulating tumor cells (CTCs) or DTCs compared to patients with IDC.^21,22^ Although this suggests dormant DTCs may drive late recurrences in patients with ILC, studies on ILC dormancy are minimal. ILC cells have low proliferative indices, resulting in few successful cell lines,^23^ patient-derived xenografts,^24^ and murine models that exhibit signs of metastasis >6 months after engraftment.^25^ Therefore, there is an unmet need to expedite ILC research while maintaining physiological relevance.

Under the United States Food and Drug Administration (FDA) Modernization Act 2.0 and 3.0, the prerequisites for human clinical trials are no longer solely based on animal models, encouraging the development of bioengineered assays for preclinical testing using human-derived cells.^26,27^ Bioengineered models have facilitated the systematic, reproducible, and high-throughput interrogation of specific cellular interactions in human-derived cells, thereby furthering our understanding of breast cancer dormancy.^28-31^ Particularly, bioengineered models have elegantly demonstrated the role of physical stimuli on cancer dormancy, including viscoelasticity,^32^ stiffness,^33-36^ and cell-binding ligand density.^37^ Given the slow-growing nature of ILC, bioengineered assays of dormancy offer a high-throughput platform to develop a comprehensive perspective on ILC dormancy.

Therefore, we employed three micro-compartmentalization techniques that encompass various dimensions of cell culture, including two-dimensional, pseudo-three-dimensional, and three-dimensional culture, enabling the investigation of cell-protein, cell-cell, and cell-matrix interactions. Inducing a dormant phenotype via oxygen and serum deprivation revealed differential dormancy patterns amongst the ILC cell lines, with p27^Kip1^ signaling at the center of the dormancy response. Interestingly, p27^Kip1^ signaling in dormant ILC cells was associated with levels of tamoxifen resistance-associated microRNA (miRNA) and chemical and mechanical sensing of the micropatterned surface. Altogether, multidimensional micro-compartmentalization enables a rapid, high-throughput method to investigate the various mechanisms governing ILC dormancy that mimic DTC dormancy.

## Results

### High-throughput cellular interactions via multidimensional micro-compartmentalization

Breast cancer cells were micro-compartmentalized utilizing three bioengineering approaches to investigate three characteristics of cancer cells. The first approach promotes the interaction of breast cancer cells with extracellular matrix (ECM) proteins on a rigid substrate through surface chemistry. Briefly, glass surfaces were coated with poly-L-lysine (PLL), which formed an amide bond with methoxy-poly(ethylene glycol)-succinimidyl valerate (mPEG-SVA) to produce a non-biofouling PLL-*graft*-mPEG (PLL-*g*-mPEG) polymer brush. The mPEG chains were spatially degraded with a photoinitiator and ultraviolet (UV) projections via a digital micromirror device (DMD), rendering the degraded regions susceptible to ECM protein adsorption and subsequent cellular attachment (**Figure 1A-i**).^38^ The method enabled two-dimensional, large-scale arrays of breast cancer cells (**Figure 1B-i**, **Supplementary Figure 1**).

**Figure 1:**
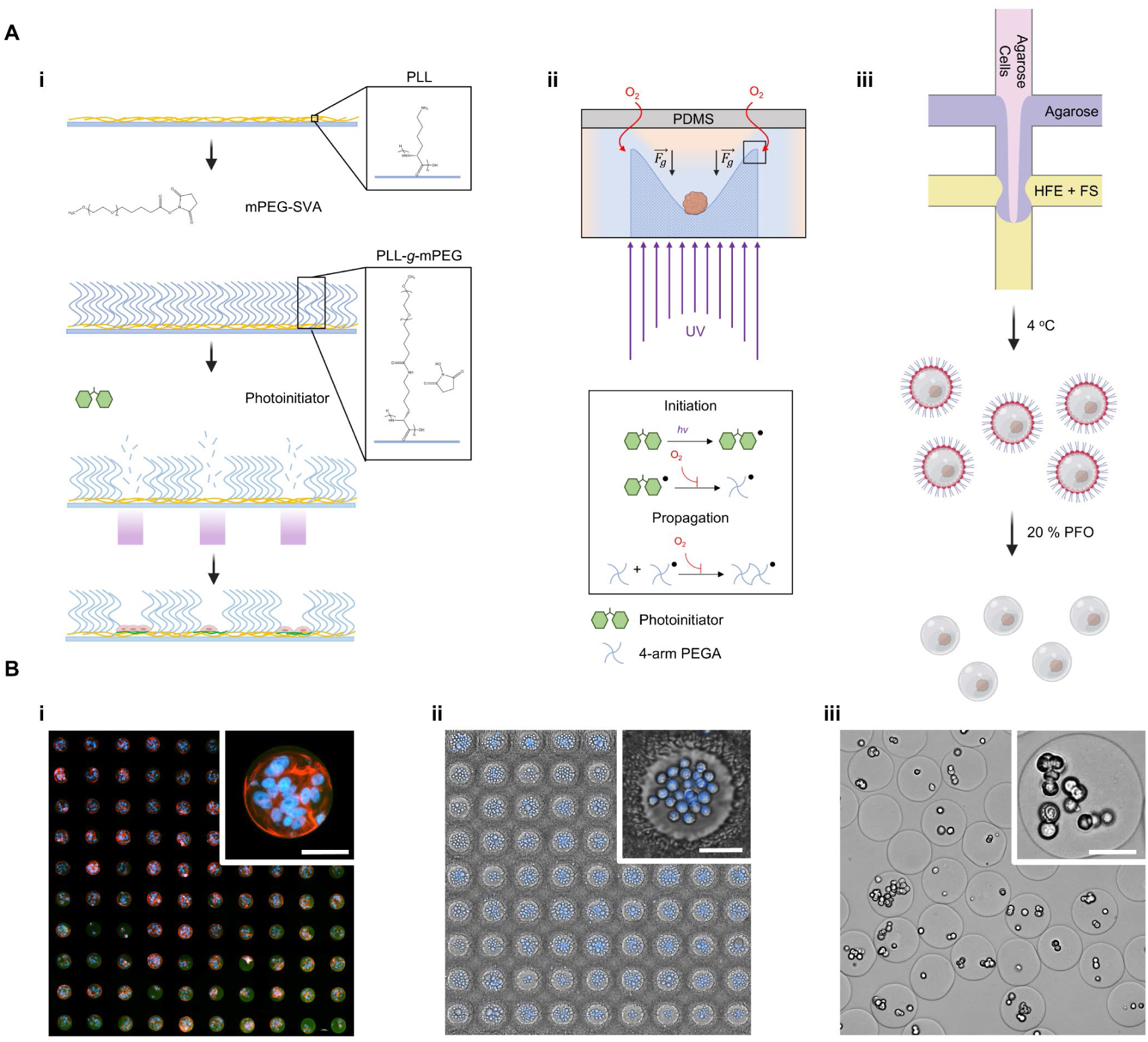
High-throughput micro-compartmentalization of breast cancer cells. (**A**) Schematic representation of the three bioengineering technologies utilized in the investigation, including (i) light-induced molecular adsorption (LIMA), (ii) microcuvettes, and (iii) microfluidic generation of microgels. (**B**) The seeding of MDA-MB-134 VI (MB134) cells within the three bioengineering technologies demonstrates the ability of (i) LIMA to isolate cell-protein interactions, (ii) the microcuvettes to induce cell-cell interaction, and (iii) the microgels to promote cell-matrix interaction. All scale bars are 50 µm.

The second technique promotes intercellular interactions within non-adhesive microcuvettes via *in situ* photopolymerization.^39^ Briefly, the DMD converted grayscale paraboloid arrays into UV projections (**Supplementary Figure 2A-B**), with grayscale values correlating to photon flux.^39^ Oxygen inhibits radical polymerization of 4-arm PEG-acrylate (4-arm PEGA) through photoinitiator quenching or chain-terminating peroxide radicals.^40^ Therefore, actuating the photon flux spatially dictated the penetration of oxygen through the poly(dimethyl siloxane) (PDMS) ceiling, realizing rapid topographical control corresponding to the grayscale image (**Supplementary Figure 2C-D**). Due to the non-adhesive nature of the PEG microcuvettes, cells aggregated under gravitational forces (**Figure 1A-ii**), allowing for the formation of cellular arrays in a pseudo-three-dimensional culture (**Figure 1B-ii**).

Lastly, the third technique promotes intercellular interaction in isotropic conditions independent of gravity and under the mechanical stress of a polymeric scaffold. Briefly, an agarose-based cell suspension was dispersed in a continuous phase comprising a fluorosurfactant (FS) diluted in hydrofluoroether (HFE) via microfluidic droplet generation.^41^ The microdroplets were crosslinked off-chip via a reduction in temperature and demulsified with 1*H*,1*H*,2*H*,2*H*-perfluoro-1-octanol (PFO; **Figure 1A-iii**). Agarose microgels were selected due to their similar mechanical properties to those of breast tissue, their optical clarity, and stability at physiological temperatures.^42^ Crosslinking and demulsifying the microdroplets facilitated a facile transition to cell growth media (**Supplementary Figure 3A-B**), with reductions in microgel size (**Supplementary Figure 3C**) and increases in the coefficient of variation, despite achieving high homogeneity (**Supplementary Figure 3D**). The physically homogeneous microgels thus enable the micro-compartmentalization of breast cancer cells in a three-dimensional culture (**Figure 1B-iii**).

### Recapitulating the morphological differences between ILC and IDC *in vitro*

MDA-MB-134 VI (MB134), MCF7, and MDA-MB-231 cells were seeded onto 100-µm fibronectin 1 (FN1) micropatterns to highlight the differences between ILC, IDC, and triple-negative breast cancer (TNBC), respectively. The MCF7 cells filled the micropatterns within 48 hours, whereas the MB134 cells remained stagnant while slowly proliferating (**Supplementary Figure 4A-B**, **Supplementary Video 1**). Interestingly, the MCF7 formed spheroidal structures, and after 2 days, extended outside the FN1 micropattern onto the PLL-*g*-mPEG surface as monolayers and expelled single cells (**Supplementary Figure 4C**, **Supplementary Video 2**). As a control, MDA-MB-231 filled the FN1 micropatterns within 1 hour and performed chiral movements along the central axis (**Supplementary Video 3**). These results indicate the variability with which different types of breast cancer cells interact with the FN1 surface. Although ILC and IDC patients receive similar treatments in the clinic, the stark differences in micropattern interaction imply significant differences between the two subtypes and warrant further comparative investigation.

To model ILC cells that persist post-treatment, MB134 and SUM44PE (SUM44) cells were made resistant to the anti-estrogen drug tamoxifen.^43^ After long-term culture with tamoxifen, the half-maximal inhibitory concentration (IC50) values for tamoxifen increased from 8.36 ± 0.34 µM to 16.38 ± 0.11 µM for the MB134 cells and 11.15 ± 0.22 µM to 27.30 ± 1.02 µM for the SUM44 cells.^43^ The parental (P) and tamoxifen-resistant (T) cells were added to the FN1 micropatterns alongside the MCF7 cells to assess differences upon acquired tamoxifen resistance. *CDH1* is deleted in ∼90% of patients with ILC,^15^ which encodes the adherence junction protein E-cadherin. As expected, only MCF7 cells expressed E-cadherin (**Figure 2A**, **Supplementary Figure 5A**). Unlike the MB134-P cells, the MB134-T cells spread across the FN1 micropattern, similar to MCF7 cells (**Figure 2A**). Given the circular design of the micropattern, the circularity and area of coverage yielded distinct population clusters that classified the cell lines. The MB134-P cells formed a distinct population with minimal circularity and area of cellular coverage (**Figure 2B**; *k*-means clustering accuracy = 98.68% for MB134-P), while MCF7 and MB134-T cells were less distinguishable from each other (**Figure 2B**; *k*-means clustering accuracy = 80.08% for MCF7, *k*-means clustering accuracy = 78.15% for MB134-T). Unlike the MB134 cells, the SUM44-T and SUM44-P cells had similar morphologies (**Supplementary Figure 5A**) with hardly discernible populations (**Supplementary Figure 5B**; *k*-means clustering accuracy = 39.47% for SUM44-P, *k*-means clustering accuracy = 75.00% for SUM44-T). After 6 days of culture, the MCF7 cells grew three-dimensionally along the *z*-axis, enriched with E-cadherin at the cell-cell junctions (**Figure 2C**). The MB134-P cells remained within the micropattern and formed single-file lines, as observed *in vivo* (**Figure 2C**).^44^ The MB134-T cells formed small three-dimensional structures of 2-3 cells in height (**Figure 2C**). E-cadherin was only present at the intercellular junctions and excluded from the extracellular domain for MCF7 cells, and was absent intercellularly and extracellularly for MB134-P and MB134-T cells (**Figure 2D**). E-cadherin aids in organizing and polarizing epithelial tissues,^45^ which may indicate the ability for MCF7 cells to form multicellular three-dimensional structures.

**Figure 2:**
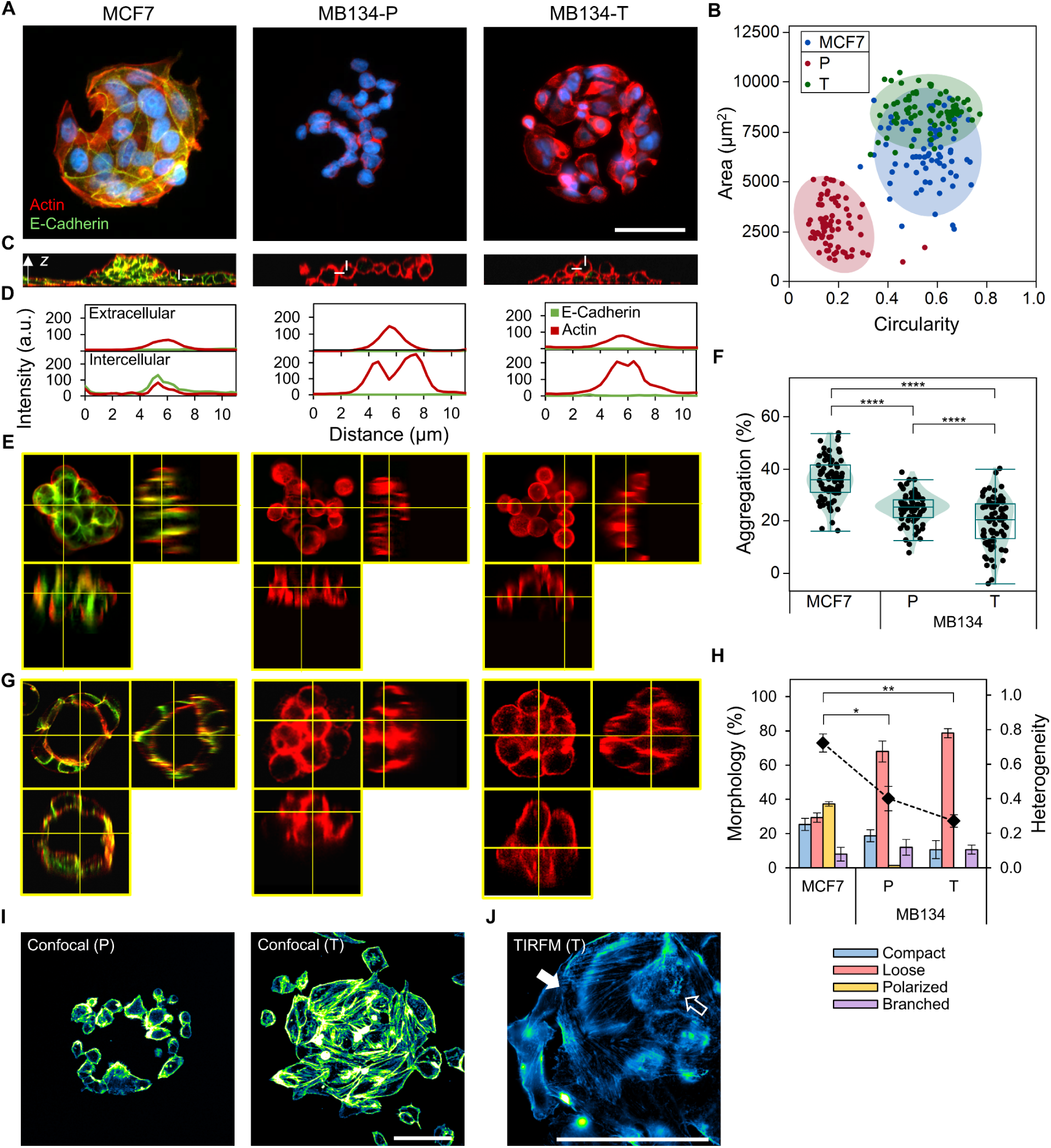
ILC and IDC cells are morphologically and molecularly distinct. (**A**) Epifluorescence microscopy of MCF7, parental MDA-MB-134 VI (MB134-P), and tamoxifen-resistant MDA-MB-134 VI (MB134-T) cells on FN1 microdomains reveals differences in interactions with the micropatterned surface and E-cadherin expression (green represents E-cadherin, red represents actin, and blue represents DAPI). (**B**) The interaction is presented as clusters of the area *versus* the circularity of cellular coverage over the microdomain (N = 3, n = 25). (**C**) The micropatterned cells form three-dimensional structures after long-term culture, as reconstructed with confocal microscopy (green represents E-cadherin and red represents actin). The white lines indicate the subsequent plane of quantification for Figure 2D. (**D**) The intercellular and extracellular expression of E-cadherin and actin in a single cell demonstrates expression of E-cadherin intracellularly only in MCF7 cells. (**E**) The confocal reconstructions of the microcuvettes indicate differences in their aggregation and organization (green represents E-cadherin and red represents actin). (**F**) The aggregation of cells, presented as violin-box plot conjugates, demonstrates differential aggregation (N = 3, n = 25; \*\*\*\**p* < 0.0001). (**G**) The confocal reconstructions of the microgels demonstrate differences in cellular organization (green represents E-cadherin and red represents actin). (**H**) Quantification of the different morphologies observed in the microgels is presented as bar graphs, and a line plot indicates the morphological heterogeneity score (N = 3, error bars indicate the standard error of the mean; \*\*\*\**p* < 0.0001). (**I**) Confocal images of cells at the coverslip surface reveal enhanced interaction and actin fibers in MB134-T cells (actin is pseudo-colored in a gradient of blue). (**J**) A TIRFM image captured at the coverslip surface shows distinct features of actin (actin is pseudo-colored in a gradient of blue). The filled arrow highlights actin polymerization, and the empty arrow highlights disorganized actin structures. All scale bars are 50 µm.

Next, all three cell lines were introduced onto the microcuvettes to determine if the differences persisted in the absence of ECM proteins. Differences in aggregation between the ILC and IDC cells were visualized at 12 hours (**Supplementary Video 4**). The MCF7 cells rapidly self-organized into compact spheroids, with E-cadherin excluded from the extracellular region and present at intercellular junctions (**Figure 2E**). The MB134 and SUM44 cells formed loose structures that were discohesive and without E-cadherin expression (**Figure 2E**, **Supplementary Figure 5C**). The extent of aggregation was highest with MCF7, followed by MB134-P cells, and was lowest for MB134-T cells (**Figure 2F**, **Supplementary Figure 6A**). To determine whether the aggregation was time-dependent, the microcuvettes were monitored for multiple days, revealing stable growth and slight increases in circularity (**Supplementary Figure 6B-C**). Again, the presence of E-cadherin seemed to facilitate the ability of MCF7 cells to form compact structures.

Lastly, the MB134 and MCF7 cells were encapsulated in microgels and allowed to reconfigure over 6 days. The MCF7 cells formed various structures, including those resembling mammary acini with intercellular E-cadherin (**Figure 2G**). On the other hand, the MB134 cells mainly formed grape-like aggregates (**Figure 2G**). Quantification of spheroid morphology revealed a heterogeneous profile for the MCF7 cells, including compact, loose, polarized, and branched structures, while the MB134 cells predominantly formed loose structures (**Figure 2H**, **Supplementary Figure 7A**). The MB134 cells continued to grow over the timeframe, becoming less compact over time (**Supplementary Figure 7B-C**). Interestingly, grape-like structures were observed in an ILC murine intraductal model.^25^ Therefore, the bioengineered techniques closely recapitulated the morphologies of ILC and IDC.

Given the distinctive morphologies of MB134-P and MB134-T cells on the FN1 patterns, the coverslip surface was visualized after 6 days of culture. Confocal microscopy revealed a stark difference in actin organization between the MB134-T cells and MB134-P cells (**Figure 2I**). Total internal reflection fluorescence microscopy (TIRFM) images of MB134-T cells revealed both the polymerization of actin as straight fibers and regions of actin disorganization on FN1 micropatterns (**Figure 2J**). Overall, the data suggest that ILC cells fail to form organized structures, whereas IDC cells form various organized structures with intercellular E-cadherin facilitating cell-cell contacts. Furthermore, significant differences were observed between the MB134-P and MB134-T cells on FN1 micropatterns but not as clearly in the microcuvettes or microgels.

### Tamoxifen-resistant ILC cells arrest at the G_0_/G_1_ phase under deprivation conditions

Based on the observation that MB134-P and MB134-T cells interact differentially with the micropatterned surface, we adsorbed two ECM proteins that affect dormancy to assess chemical cues, including FN1, which promotes proliferation,^46^ and collagen type III, alpha 1 (COL3A1), which promotes dormancy.^47^ Interestingly, the MB134-P cells demonstrated preferential attachment to COL3A1, as opposed to the MB134-T cells that preferentially bound to FN1 (**Figure 3A**). Serum and oxygen deprivation can independently induce a dormant cellular phenotype,^48-50^ but were performed in conjunction for 48 hours to induce dormancy in the ILC cells. The ILC cells were transformed with a PIP-mVenus-P2A-mCherry-Gem_1-110_ (PIP-FUCCI) biosensor to track the cell cycle, where cells in the G_0_/G_1_ phase fluoresce green, in the S phase fluoresce red, and in the G2/M phase fluoresce yellow (**Figure 3B**, **Supplementary Figure 8**, **Supplementary Video 5**).^51^ To determine whether the effects of deprivation were not detrimental to the cell, we measured viability across the different bioengineered platforms. The viability was maintained above 80% except for the microgels under deprivation conditions, with viabilities of 76.01% for MB134-P and 40.73% for MB134-T (**Figure 3C**). Although the viabilities in the microgel were low, after 6 days of culture under deprivation conditions, the MB134-T cells were unable to form spheroids, whereas the MB134-P cells were less perturbed by the insult (**Supplementary Figure 9A-C**). The distributions of cells per microgel shifted to the left for the MB134-T cells, likely due to cell death, and moved to the right for the MB134-P cells, likely due to minimal growth (**Supplementary Figure 9D-F**). The mechanical stress imposed by the bioinert matrix, in addition to the stress induced by serum and oxygen deprivation, appeared to have a cumulative effect on the MB134 cells, potentially affecting dormancy readouts.

**Figure 3:**
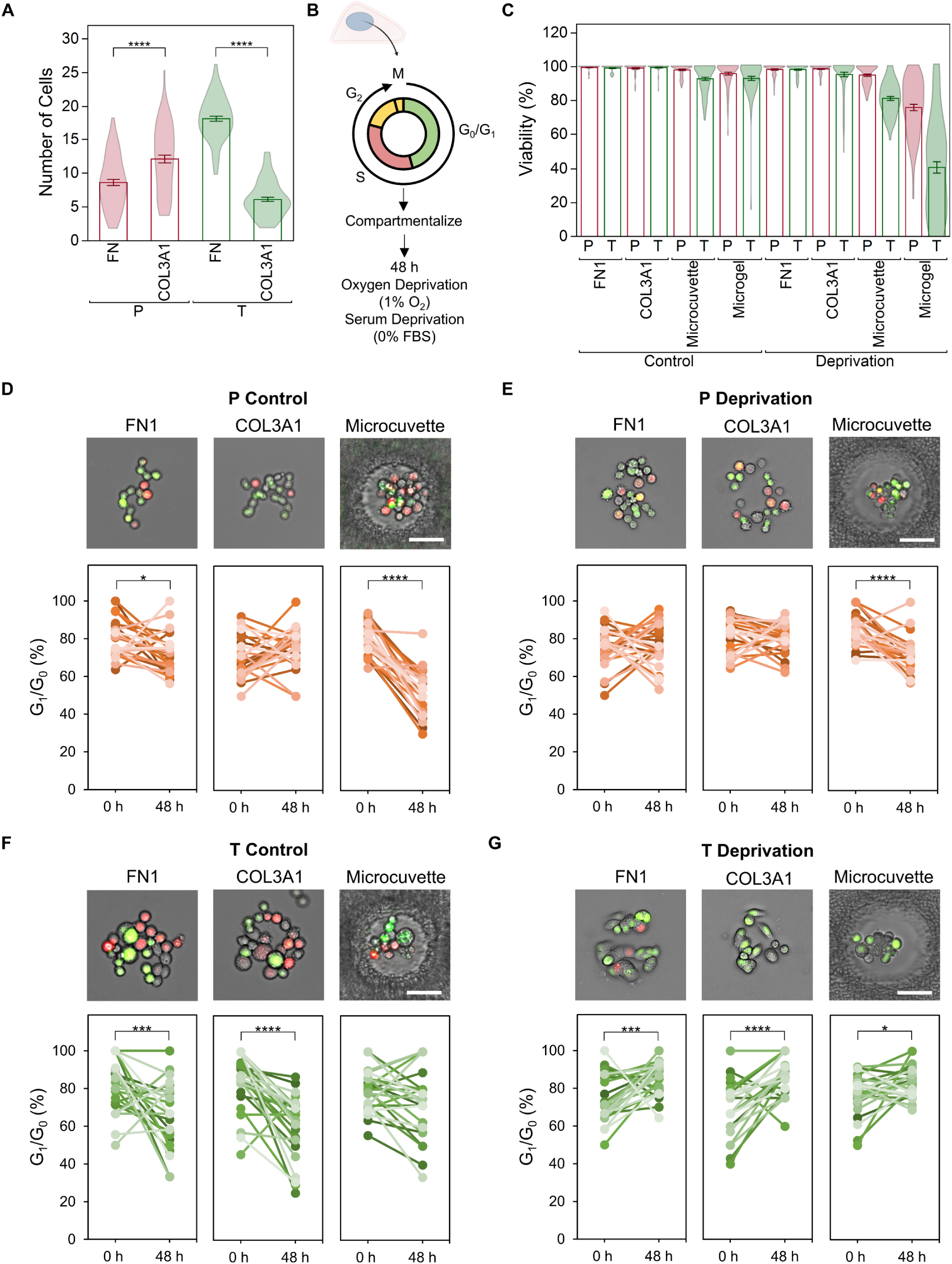
MB134-T cells enter the G_1_/G_0_ phase more than MB134-P cells. (**A**) The number of cells attached for MB134-P and MB134-T cells is different for FN1 and COL3A1 micropatterns (N = 3, n = 25, error bars indicate the standard error of the mean; \*\*\*\**p* < 0.0001). (**B**) Schematic of the cell-cycle biosensor and the protocol for deprivation-induced dormancy. (**C**) Viability across the bioengineered techniques reveals cell death with the microgels under deprivation conditions (N = 3, n = 25, error bars indicate the standard error of the mean). (**D**) Representative images at 48 hr and the corresponding quantification of the live-cell tracking of the biosensor in MB134-P cells under control conditions across bioengineered techniques (n = 25; \**p* < 0.05, \*\*\*\**p* < 0.0001). (**E**) Representative images at 48 hr and the corresponding quantification of the live-cell tracking of the biosensor in MB134-P cells under deprivation conditions across bioengineered techniques (n = 25; \*\*\*\**p* < 0.0001). (**F**) Representative images at 48 hr and the corresponding quantification of the live-cell tracking of the biosensor in MB134-T cells under control conditions across bioengineered techniques (n = 25; \*\*\**p* < 0.001, \*\*\*\**p* < 0.0001). (**G**) Representative images at 48 hr and the corresponding quantification of the live-cell tracking of the biosensor in MB134-T cells under deprivation conditions across bioengineered techniques (n = 25; \**p* < 0.05, \*\*\**p* < 0.001, \*\*\*\**p* < 0.0001). All scale bars are 50 µm.

Therefore, we advanced the investigation using only micropatterns and microcuvettes to control for cell death as a covariate. Following the same micro-compartmentalized MB134 cells over 48 hours, significant loss of cells in the G_0_/G_1_ phase was observed for the MB134-P cells on FN1 micropatterns and microcuvettes, but not on COL3A1 micropatterns under control conditions (**Figure 3D**). Under deprivation conditions, the loss of the MB134-P cells in the G_0_/G_1_ phase was reduced and did not reach significance, while the microcuvette exhibited a significant loss (**Figure 3E**). Similar to the MB134-P cells, the MB134-T cells exhibited significant cell loss in the G_0_/G_1_ phase, but on both FN1 and COL3A1, and not in the microcuvettes (**Figure 3F**). Interestingly, under deprivation conditions, the MB134-T cells demonstrated enhanced accumulation in the G_0_/G_1_ phase on all platforms, unlike the MB134-P cells (**Figure 3G**). Therefore, the MB134-T cells appear to have an enhanced sensitivity to the deprivation condition, entering the G_0_/G_1_ phase, indicative of dormancy, which was less pronounced for the MB134-P cells.

### ILC and IDC cells differentially enter dormant phenotypes

To corroborate the transition into the G_0_/G_1_ phase of the cell cycle with dormancy, we investigated several markers implicated in cancer dormancy, quiescence, senescence, and proliferation on the FN1 and COL3A1 micropatterns. The cyclin-dependent kinase (CDK) inhibitor p27^Kip1^ was dramatically increased under deprivation conditions in both MB134-P and MB134-T cells (**Figure 4A**) and was present in the majority of MB134-T cells on COL3A1 micropatterns (**Supplementary Figure 10A**). Further investigation of the deprivation conditions revealed that p27^Kip1^ was upregulated to a greater extent in MB134-T cells compared to MB134-P cells, and depended on the micropatterned protein. Specifically, p27^Kip1^ expression increased almost twofold in MB134-T cells on COL3A1 compared to FN1 micropatterns, whereas it remained unchanged in MB134-P cells (**Figure 4A**). Interestingly, the proliferation marker, Ki-67, did not follow the same trend. The MB134-T cells exhibited overall higher expression than the MB134-P cells, with a higher expression on FN1 micropatterns compared to COL3A1 micropatterns for both cell lines. A slight reduction in Ki-67 expression under deprivation conditions was observed for the MB134-P cells, but not for the MB134-T cells (**Supplementary Figure 10B**). The data suggest that a subpopulation of cells undergoes cell cycle arrest, while others continue to proliferate. To determine the role of another compensatory pathway, the CDK inhibitor p21^Cip1/Waf1^ was screened, revealing a decrease in expression for the MB134-P cells and no change for the MB134-T cells (**Supplementary Figure 10C**). Since the proliferative indices varied between the MB134-P and MB134-T cells, Ki-67 and p27^Kip1^ were screened over longer timeframes on COL3A1 micropatterns. Although Ki-67 expression decreased in both MB134-T and MB134-P cells over 6 days under deprivation conditions, the expression of Ki-67 drastically reduced in the MB134-T cells after 2 days under control conditions, due to the confluence of the micropatterned surface (**Supplementary Figure 10D**). In contrast, the expression of Ki-67 in MB134-P cells initially increased and declined after 4 days (**Supplementary Figure 10D**). On the other hand, p27^Kip1^ expression continually increased in the MB134-T cells under both deprivation and control conditions, the latter being influenced by the confluence of the micropatterned surface (**Supplementary Figure 10E**). In contrast, the MB134-P cells showed no change in p27^Kip1^ expression (**Supplementary Figure 10E**). Furthermore, a comparison of 200-µm micropatterns *versus* 100-µm micropatterns revealed similar profiles with slight decreases in Ki-67 and p21^Cip1/Waf1^ but no change in p27^Kip1^ expression under deprivation conditions (**Supplementary Figure 10F**), indicating that cell confluency is not driving quiescence under deprivation conditions. These results suggest that 48 hours post-seeding provides the most significant differences between the control and deprivation conditions, controlling for confluency as a covariate. Therefore, the differences in arrest at the G_1_/G_0_ cycle between MB134-P and MB134-T cells are mediated by p27^Kip1^ signaling, with MB134-T cells demonstrating sensitivity to the chemical cues of the micropatterned surface.

**Figure 4:**
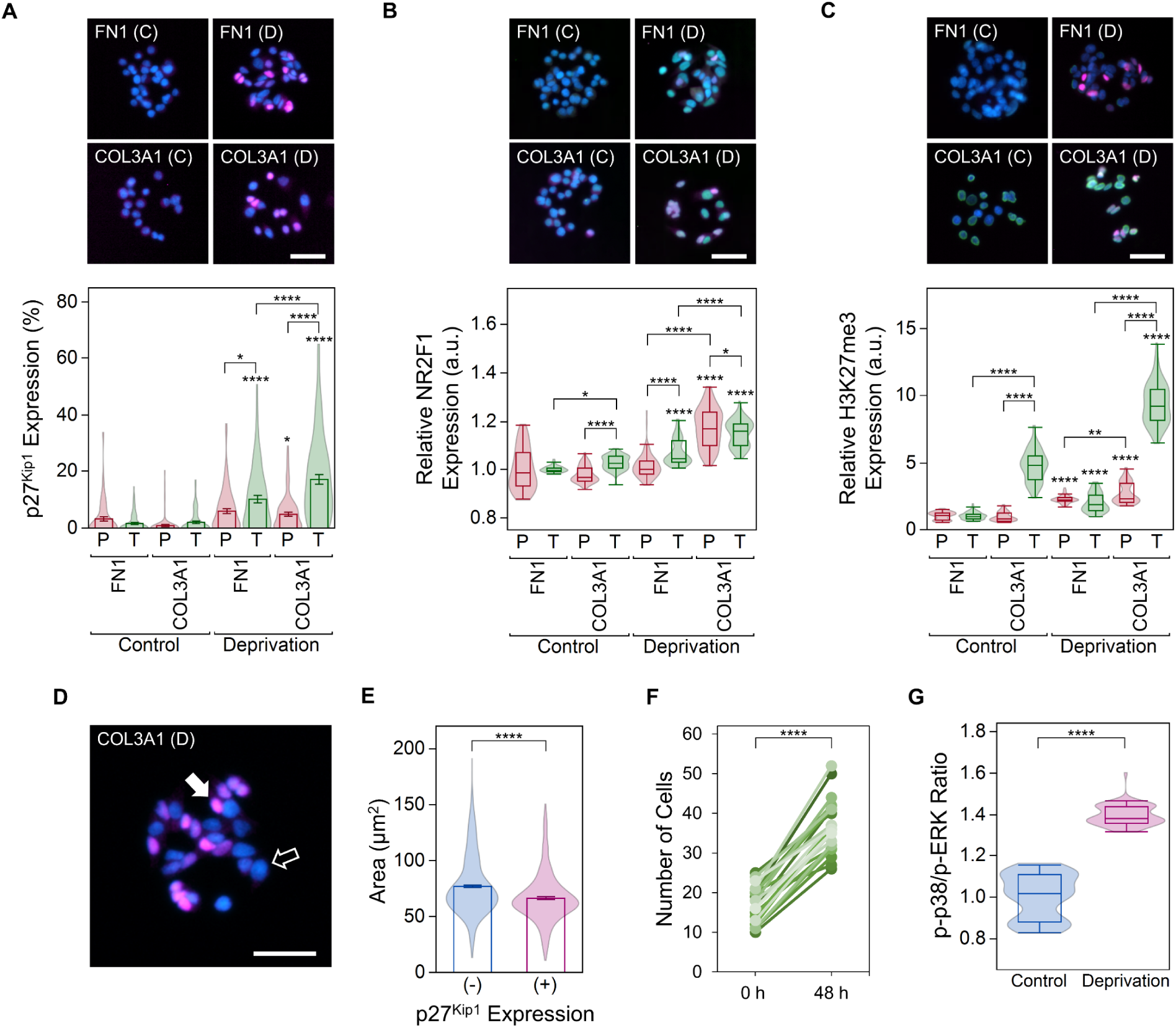
MB134 cells undergo p27^Kip1^-mediated dormancy. (**A**) Representative epifluorescence microscopy of MB134-T cells (magenta represents p27^Kip1^ and blue represents DAPI) is supplemented by the expression profiles of p27^Kip1^ as violin-bar conjugate plots, demonstrating the highest expression for the MB134-T cells on COL3A1 under deprivation condition (N = 3, n = 25, error bars indicate the standard error of the mean; \**p* < 0.05, \*\*\*\**p* < 0.0001). (**B**) Representative epifluorescence microscopy of MB134-T cells (magenta represents p27^Kip1^, NR2F1 represents green, and blue represents DAPI) is supplemented by the expression profiles of NR2F1 as violin-box conjugate plots, demonstrating protein-dependent expression for the MB134-P and MB134-T cells under deprivation conditions (N = 3, n = 25; \**p* < 0.05, \*\*\*\**p* < 0.0001). (**C**) Representative epifluorescence microscopy of MB134-T cells (magenta represents p27^Kip1^, H3K27me3 represents green, and blue represents DAPI) is supplemented by the expression profiles of H3K27me3 as violin-box conjugate plots, demonstrating protein-dependent expression that is augmented by the MB134-T cells under deprivation conditions (N = 3, n = 25; \*\**p* < 0.0001, \*\*\*\**p* < 0.0001). (**D**) Representative epifluorescence microscopy of the MB134-T cells on a COL3A1 microdomain illustrates that p27^Kip1^-expressing cells have smaller nuclei (filled arrow) compared to p27^Kip1^-negative cells that have larger nuclei (empty arrow; magenta represents p27^Kip1^ and blue represents DAPI). (**E**) Quantification of nuclear size for MB134-T cells on COL3A1 micropatterns in deprivation conditions, presented as violin-bar conjugate plots, further demonstrates the difference in size (N = 3, n = 25, error bars indicate the standard error of the mean; \*\*\*\**p* < 0.0001). (**F**) Upon reintroduction to control conditions for 48 hr, the deprived MB134-T cells proliferate, exhibiting the reversibility of the deprivation-induced dormant phenotype (n = 25, error bars indicate the standard error of the mean; \*\*\*\**p* < 0.0001). (**G**) The p-p38/p-ERK ratio, represented as violin-box conjugate plots, is elevated for MB134-T cells under deprivation conditions (N = 3, n = 25; \*\*\*\**p* < 0.0001). All scale bars are 50 µm.

To further correlate the observed p27^Kip1^ signaling with DTC dormancy, we screened for nuclear receptor subfamily 2, group F, member 1 (NR2F1), a marker that is upregulated in dormant DTCs *in vivo*.^48,52-54^ Similar to p27^Kip1^, the micropatterned proteins and conditions influenced NR2F1 expression, where COL3A1 compared to FN1 micropatterns led to higher expression for both MB134 cells under deprivation conditions (**Figure 4B**). Furthermore, H3K27me3 was screened since chromatin states in dormant DTCs are altered downstream of NR2F1.^53^ Although the H3K27me3 protein was increased in both MB134 cells under deprivation conditions, the expression was significantly enhanced in MB134-T cells on COL3A1 micropatterns (**Figure 4C**). These findings confirm that the differences in the transition to the G_1_/G_0_ cycle and p27^Kip1^ signaling portrayed by our system recapitulate a similar dormant phenotype observed in DTCs *in vivo*.^48,52-54^

Cells escape the cell cycle through various mechanisms. Dormancy is often associated with quiescence, a reversible form of cell cycle arrest, as opposed to senescence, which is an irreversible form of cell cycle arrest. Therefore, we aimed to distinguish quiescence from senescence. Nuclear size is often a descriptor of cell-cycle arrest, in which quiescent cells display smaller nuclei,^55^ and senescent cells exhibit larger nuclei.^56,57^ Interestingly, p27^Kip1^-expressing cells presented smaller nuclei compared to non-expressing cells on COL3A1 micropatterns under deprivation conditions (**Figure 4D-E**). To determine the reversibility of the dormant phenotype, cells were incubated under deprivation conditions for 48 hours on COL3A1 micropatterns and then returned to control conditions for an additional 48 hours. All cells on the micropatterns continued their growth after transition (**Figure 4F**). As expected, Ki-67 expression increased upon restoration of control conditions (**Supplementary Figure 11A**) and decreased p27^Kip1^ substantially (**Supplementary Figure 11B**). Furthermore, CDK inhibitors associated with senescence, p21^Cip1/Waf1^ and p16^Ink4a^, were unchanged in the MB134-T cells under deprivation conditions (**Supplementary Figure 10C**, **Supplementary Figure 11C**). Lastly, the p-p38/p-ERK ratio, associated with reversible dormancy,^46^ was elevated in MB134-T cells on COL3A1 micropatterns under deprivation conditions (**Figure 4G**). These results demonstrate an ability to replicate a reversible dormant phenotype with clinical relevance to late recurrences.

To test whether the observed signaling in MB134-T cells was specific to deprivation-induced dormancy, we used interferon (IFN)-γ to simulate immunologic stress and induce dormancy.^58^ While there was a slight but significant increase in Ki-67 with IFN-γ treatment, irrespective of the micropatterned protein (**Supplementary Figure 12A**), both p21^Cip1/Waf1^ and p27^Kip1^ were significantly increased with IFN-γ treatment, where COL3A1 compared to FN1 micropatterns induced relatively higher expression of both CDK inhibitors (**Supplementary Figure 12B-C**). Interestingly, both dormancy-inducing methods yielded a protein-dependent expression profile of p27^Kip1^ in the MB134-T cells. However, given the increase in p21^Cip1/Waf1^ following IFN-γ treatment, we advanced our studies by examining deprivation-induced dormancy to control for senescence as a covariate.

Unlike the MB134 cells, the SUM44-P cells preferentially adhered to FN1, while the SUM44-T cells demonstrated no preference (**Supplementary Figure 13A**). Similar to the MB134 cells, the SUM44-T cells had higher Ki-67 expression than SUM44-P cells, except for COL3A1 micropatterns under control conditions (**Supplementary Figure 13B**). Furthermore, there was a decrease in Ki-67 expression under deprivation conditions for SUM44-P cells on COL3A1 micropatterns and SUM44-T cells on FN1 micropatterns (**Supplementary Figure 13B**). While similar to the MB134 cells, p21^Cip1/Waf1^ was most highly expressed in SUM44-P cells but was unaffected under deprivation conditions in both SUM44 cells (**Supplementary Figure 13C**). Furthermore, the SUM44 cells exhibited a broad elevation in p27^Kip1^ expression post-deprivation but without sensitivity to the micropatterned protein or differences in expression between the SUM44-T and SUM44-P cells (**Supplementary Figure 13D**). The expression of NR2F1 in SUM44-T was broadly elevated post-deprivation; however, unlike the MB134-T cells, the interaction between the condition and the micropatterned protein was absent (**Supplementary Figure 13E**). Similarly, the SUM44-T upregulated H3K27me3 under deprivation conditions without interaction between the condition and the micropatterned protein (**Supplementary Figure 13F**). Interestingly, both MB134-T and SUM44-T cells under the control conditions demonstrated a spatial expression of H3K27me3, albeit at lower expression levels compared to the deprivation conditions (**Supplementary Figure 14**). The spatial expression resembles a phenomenon observed in migratory leader cells in epithelial tissues.^59^

Lastly, to compare fast-cycling IDC cells with slow-cycling ILC cells,^60,61^ MCF7 cells were cultured on the FN1 and COL3A1 micropatterns. Indeed, the MCF7 cells showed significantly higher proliferative indices compared to the MB134-P and MB134-T cells (**Supplementary Figure 15A**). Similar to the MB134-T cells, the MCF7 cells preferentially adhered to FN1 (**Supplementary Figure 15B**), which corresponds with the finding that fast-cycling cancer cells preferentially adhere to FN1.^46^ Unlike the MB134 cells, Ki-67 was downregulated in the MCF7 cells under the deprivation conditions, irrespective of the micropatterned protein (**Supplementary Figure 15C**). While p21^Cip1/Waf1^ expression was high in MCF7 cells under control conditions, the deprivation conditions had no effect (**Supplementary Figure 15D**). Coinciding with the MB134 and SUM44 cells, p27^Kip1^ was upregulated in the MCF7 cells under deprivation conditions, albeit independent of the micropatterned protein (**Supplementary Figure 15E**). Interestingly, the MCF7 cells transitioned to a classical dormant state, characterized by a decrease in Ki-67 and an increase in p27^Kip1^. In contrast, the MB134 and SUM44 cells clearly upregulated p27^Kip1^ but yielded less defined Ki-67 profiles, highlighting the importance of treating IDC and ILC as separate entities.

### Tamoxifen resistance-associated miRNA correlates with p27^Kip1^ expression

We observed differences in the expression of p27^Kip1^ under deprivation conditions, particularly on COL3A1, that led us to investigate the molecular differences between the four ILC cell lines. Our previous studies showed that hsa-miR-221-3p and hsa-miR-222-3p confer tamoxifen resistance in IDC cells and are upregulated in HER2/neu-negative primary breast cancer samples.^62^ Screening of these miRNAs revealed the upregulation of hsa-miR-221-3p and hsa-miR-222-3p in the MB134-T cells compared to the MB134-P cells, which was not observed for SUM44 cells (**Figure 5A**). SUM44 cells were derived from the pleural effusion of a patient with ILC unresponsive to both anti-estrogen therapy and chemotherapy,^63^ unlike MB134 cells, which were derived from the pleural effusion of a patient with ILC responsive to chemotherapy.^64^

**Figure 5:**
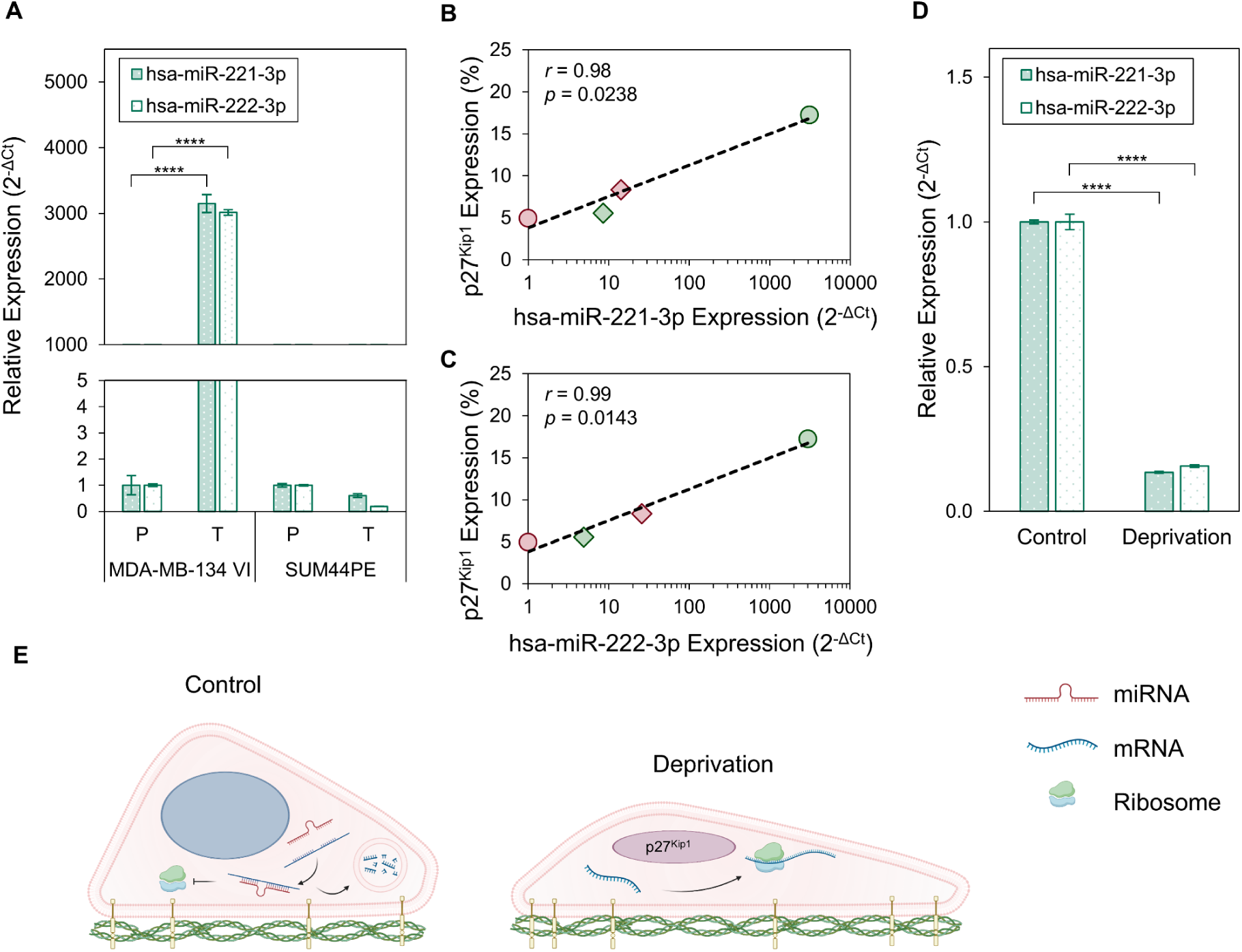
Correlation of tamoxifen resistance-associated miRNA and p27^Kip1^. (**A**) The relative expression of hsa-miR-221-3p and hsa-miR-222-3p (relative to MB134-P and SUM44-P) indicates an increased expression of the miRNA in the MB134-T cells (N = 3, error bars indicate the standard error of the mean; \*\*\*\**p* < 0.0001). (**B**) The expression of p27^Kip1^ on COL3A1 micropatterns correlates with the expression of hsa-miR-221-3p. (**C**) The expression of p27^Kip1^ on COL3A1 micropatterns correlates with the expression of hsa-miR-222-3p. (**D**) The relative expression of hsa-miR-221-3p and hsa-miR-222-3p (relative to the control conditions) reveals a loss of the miRNA under deprivation conditions (N = 3, error bars indicate the standard error of the mean; \*\*\*\**p* < 0.0001). (**E**) A schematic representation of MB134-T cells shows that under control conditions, increases in hsa-miR-221-3p and hsa-miR-222-3p are active, blocking the translation of p27^Kip1^; whereas, deprivation conditions enable the production of p27^Kip1^ due to reduced miRNA levels.

Interestingly, hsa-miR-221-3p and hsa-miR-222-3p are well-known post-translational suppressors of p27^Kip1^ by binding to two sites of the 3’-untranslated region (3’UTR) of the encoding mRNA.^65^ Surprisingly, the expression of p27^Kip1^ post-deprivation on COL3A1 micropatterns correlated with the logarithmic native levels of hsa-miR-221-3p and hsa-miR-222-3p in the four cell lines as visualized by a scatter plot (**Figure 5B-C**). To investigate the correlation between tamoxifen resistance-associated miRNA, which is highly upregulated in MB134-T cells, and their enhanced expression of p27^Kip1^, the perturbation of hsa-miR-221-3p and hsa-miR-222-3p expression post-deprivation was evaluated in MB134-T cells. Indeed, hsa-miR-221-3p and hsa-miR-222-3p were downregulated more than fivefold under the deprivation conditions (**Figure 5D**). Therefore, MB134-T cells appear to undergo epigenetic modifications via miRNA downregulation and NR2F1-mediated chromatin repression, which induce p27^Kip1^-mediated dormancy, where the former delineates the differences observed between the four ILC cell lines (**Figure 5E**).

### Dormancy-inducing conditions lead to a mesenchymal-like phenotype

Dormant DTCs acquire a mesenchymal-like phenotype.^66,67^ Therefore, we investigated the effects of the deprivation conditions on the morphology of MB134-T cells on COL3A1 micropatterns. The MB134-T cells on the COL3A1 micropatterns had rounder morphologies under control conditions, whereas deprivation conditions induced cell elongation (**Figure 6A**). To further clarify the morphology of the dormant MB134-T cells, the cytoskeletal proteins actin and β-tubulin were visualized using TIRFM. The TIRFM images revealed actin-rich protrusions reminiscent of filopodia (**Figure 6B**). Scanning electron microscopy (SEM) further confirmed the organization of fibrils in the lamellipodia that extend toward filopodia in deprived MB134-T cells, otherwise absent under the control conditions (**Figure 6C**) or in deprived MB134-P cells (**Supplementary Figure 16**). Actin-rich protrusions are often utilized in chemical sensing, as exemplified by phagocytic immune cells,^68,69^ which may further explain the sensitivity of MB134-T but not MB134-P cells or the SUM44 cells to the micropatterned protein. Furthermore, higher aspect ratios are observed for mesenchymal-like DTCs,^67^ which was recapitulated by the MB134-T cells on the COL3A1 micropatterns under deprivation conditions (**Figure 6D**). Therefore, the deprivation conditions appear to induce a mesenchymal-like morphology in the MB134-T cells.

**Figure 6:**
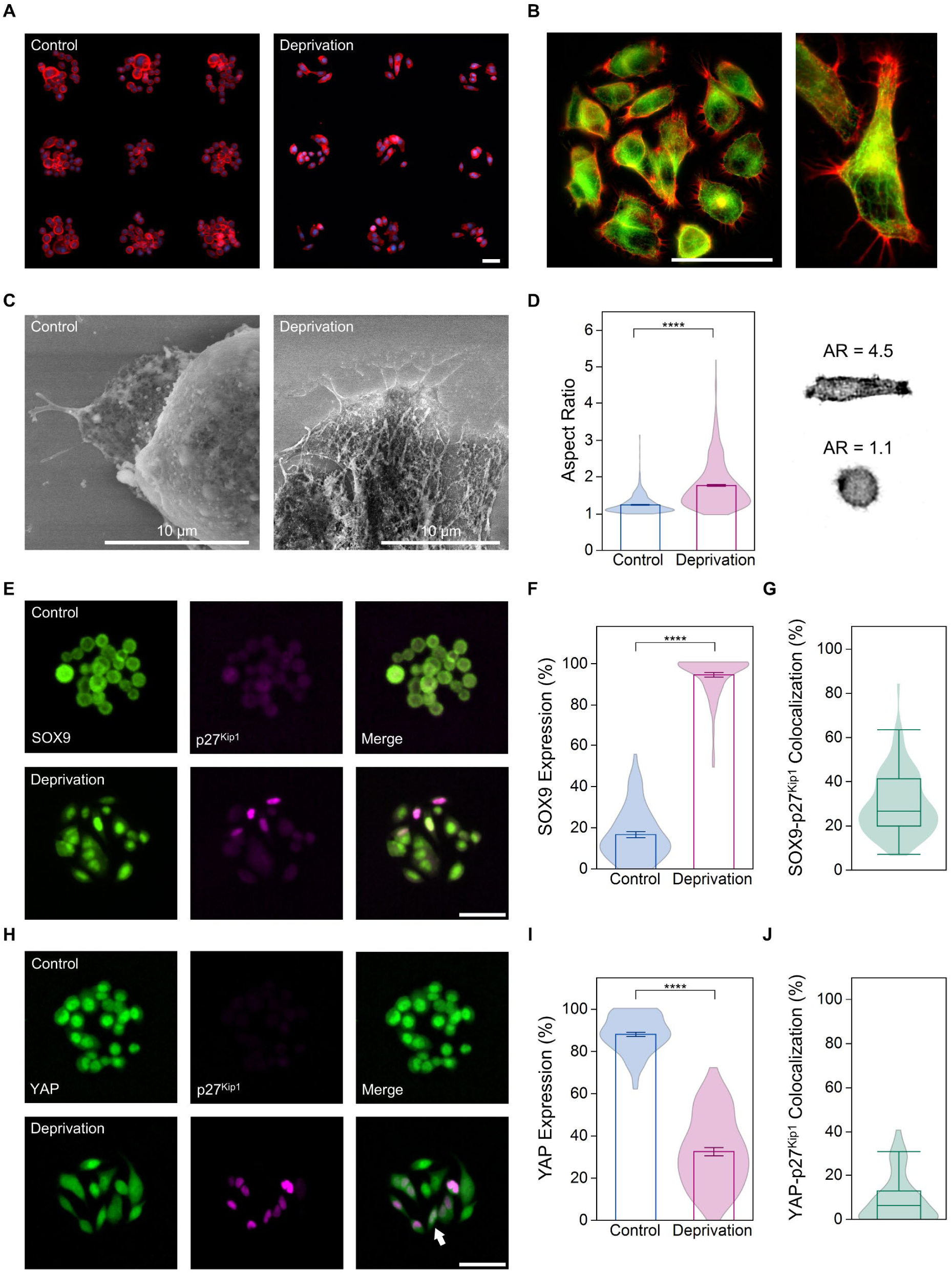
Deprived MB134-T cells exhibit a mesenchymal-like and mechanosensing phenotype. (**A**) Epifluorescence microscopy of a 3 × 3 array of MB134-T cells on COL3A1 micropatterns subjected to control and deprivation conditions shows an elongated morphology under the deprivation conditions (red represents actin and blue represents DAPI). (**B**) A TIRFM image of cells at the coverslip surface shows the formation of actin-rich filopodia in MB134-T cells on COL3A1 micropatterns under the deprivation condition (red represents actin and green represents β-tubulin). (**C**) SEM images of the MB134-T cells on COL3A1 micropatterns subject to the deprivation condition confirm the presence of tubular structures extending toward filopodia. (**D**) The aspect ratio presented in the violin-bar conjugate plots demonstrates that MB134-T cells are more elongated when subject to deprivation conditions (N = 3, n = 25, error bars indicate the standard error of the mean; \*\*\*\**p* < 0.0001). The adjacent images illustrate the morphologies associated with different aspect ratios. (**E**) Representative epifluorescence microscopy of SOX9 and p27^Kip1^ expression in MB134-T cells exhibits the nuclear translocation of SOX9 under deprivation conditions. (**F**) The nuclear SOX9 level is quantified in violin-bar conjugate plots (N = 3, n = 25, error bars indicate the standard error of the mean; \*\*\*\**p* < 0.0001). (**G**) The nuclear colocalization of SOX9 and p27^Kip1^ under the deprivation condition is quantified in a violin-box conjugate plot (N = 3, n = 25). (**H**) Representative epifluorescence microscopy of YAP and p27^Kip1^ expression exhibits the cytoplasmic translocation of YAP in MB134-T cells under deprivation conditions. (**I**) The nuclear expression of YAP is quantified in the violin-bar conjugate plots (N = 3, n = 25, error bars indicate the standard error of the mean; \*\*\*\**p* < 0.0001). (**J**) The nuclear colocalization of YAP and p27^Kip1^ under the deprivation condition is presented in a violin-box conjugate plot (N = 3, n = 25). All scale bars are 50 µm unless otherwise indicated.

To further confirm the mesenchymal-like morphology as a phenotype, we screened for SOX9 expression, a transcription factor responsible for maintaining progenitor pools throughout embryonic development.^70^ Similarly, tumor cells are programmed to acquire stem cell-like properties through SOX9, which is expressed in CTCs and DTCs that are prone to dormancy.^52,53^ Under deprivation conditions, SOX9 translocated from the cytoplasm to the nucleus in the MB134-T cells (**Figure 6E-F**), with a modest colocalization of p27^Kip1^ and SOX9 (**Figure 6G**). In summary, dormant MB134-T cells undergo a mesenchymal-like reprogramming.

### Removal of the protein-cell interactions promotes a deeper quiescence phenotype

Apart from chemical sensing, filopodia are implicated in mechanosensing.^71-73^ We screened the MB134-T cells for the yes-associated protein (YAP), whose nuclear translocation is associated with mechanosensing and subsequent signal transduction.^74^ As expected, under the control condition, MB134-T cells on the COL3A1 micropatterned rigid glass surface showed the presence of YAP predominantly in the nucleus (**Figure 6H**). However, under deprivation conditions, the nuclear localization of YAP was markedly reduced, especially in the cells positive for p27^Kip1^ (**Figure 6H-I**). Notably, the nuclear colocalization of p27^Kip1^ and YAP in cells under deprivation conditions was low (**Figure 6J**). YAP is central to the Hippo pathway and known to influence proliferation,^75-77^ suggesting that mechanosensing may be responsible for the subpopulation of proliferative cells previously observed.

To eliminate cellular interaction with the micropattern and limit mechanosensing, we employed two approaches: physical inhibition using microcuvettes and pharmacological inhibition with a focal adhesion kinase (FAK) and Pyk2 inhibitor, PND-1186 (**Figure 7A**). Indeed, when plated on the microcuvettes, the MB134-T cells excluded YAP from the nucleus, which was predominantly present in the cytoplasm, as visualized by a confocal reconstruction of the microcuvettes (**Figure 7B**). Ki-67 expression for both MB134-T and MB134-P cells on the microcuvettes was significantly reduced under deprivation conditions, with the MB134-T cells losing almost all expression (**Figure 7C**). The extent of Ki-67 downregulation in MB134-T cells was significantly greater than that in MB134-P cells under deprivation conditions (**Figure 7D**). c, p27^Kip1^ was not significantly increased on the microcuvettes compared to COL3A1 but did yield a deprivation- and cell-line-dependent profile as previously observed (**Figure 7E**). Similarly, p21^Cip1/Waf1^ was unaffected by the microcuvette and exhibited deprivation-dependent decreases, as previously observed (**Supplementary Figure 17**). Notably, MCF7 also experienced a loss of Ki-67 expression on the microcuvettes, slight increases in p21^Cip1/Waf1^, and substantial increases in p27^Kip1^ (**Supplementary Figure 18A-C**). These results indicate that the MB134 cells undergo a more classical dormant state in the microcuvettes, similar to that of MCF7 cells on the micropatterned surface or the microcuvettes. Therefore, chemical and mechanical sensing influence the dormancy phenotype of MB134 cells, suggesting that microcuvettes are the most suitable model for screening the effects of agents on ILC dormancy.

**Figure 7:**
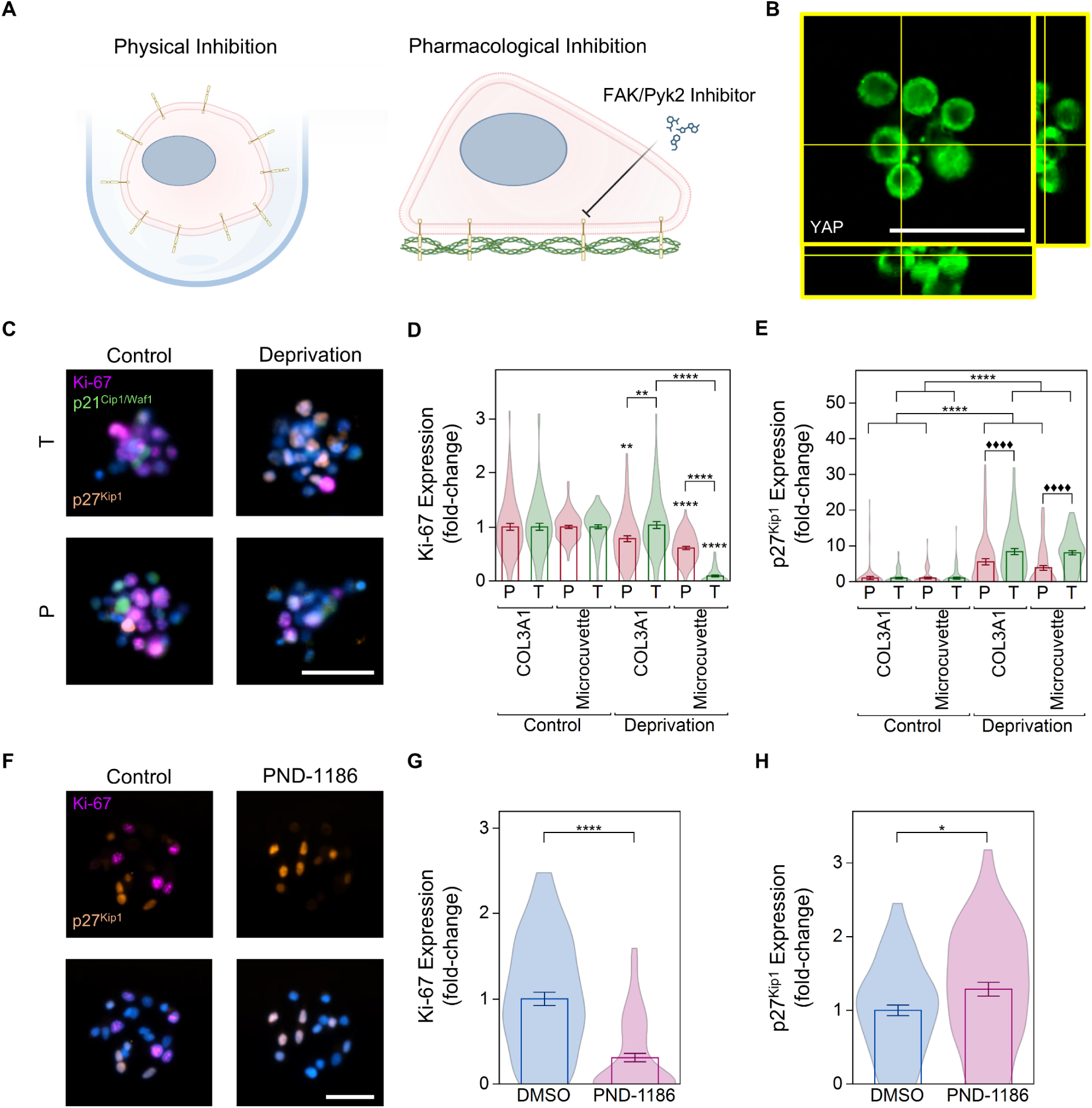
Inhibition of cell-protein interactions reduces Ki-67 in MB134-T cells. (**A**) A schematic representation of the methods to inhibit cell-protein interactions via physical and pharmacological inhibition. (**B**) The confocal reconstruction of a microcuvette reveals that YAP is predominantly located in the cytoplasm. (**C**) Representative epifluorescence microscopy of MB134-T and MB134-P cells (magenta represents Ki-67, green represents p21^Cip1/Waf1^, orange represents p27^Kip1^, and blue represents DAPI) demonstrates a loss of Ki-67 and the upregulation of p27^Kip1^ under deprivation conditions, particularly for MB134-T cells. (**D**) The expression of Ki-67 comparing COL3A1 micropatterns and microcuvettes is reduced under deprivation conditions, especially on the microcuvettes (N = 3, n = 25, error bars indicate the standard error of the mean; \*\**p* < 0.01, \*\*\*\**p* < 0.0001). (**E**) The expression of p27^Kip1^ is upregulated under deprivation conditions, with microcuvettes and COL3A1 micropatterns revealing similar levels of upregulation (N = 3, n = 25, error bars indicate the standard error of the mean; \*\**p* < 0.01, \*\*\*\**p* < 0.0001, ^♦♦♦♦^*p* < 0.0001 for the group). (**F**) Representative epifluorescence microscopy of MB134-T under PND-1186 treatment (magenta represents Ki-67, orange represents p27^Kip1^, and blue represents DAPI) demonstrates a loss of Ki-67 and the upregulation of p27^Kip1^ under treatment conditions. (**G**) The expression of Ki-67, represented as violin-bar conjugate plots, is reduced under treatment conditions (N = 3, n = 25, error bars indicate the standard error of the mean; \*\*\*\**p* < 0.0001). (**H**) The expression of p27^Kip1^, represented as violin-bar conjugate plots, is slightly upregulated under treatment conditions (N = 3, n = 25, error bars indicate the standard error of the mean; \**p* < 0.05). All scale bars are 50 µm.

Interestingly, a similar phenotype was observed in MB134-T cells on COL3A1 micropatterns when FAK/Pyk2 was pharmacologically inhibited using PND-1186, resulting in a reduction of Ki-67 (**Figure 7F**). PND-1186 treatment markedly reduced Ki-67 expression compared to the empty vehicle (**Figure 7G**) and demonstrated a small but significant increase in p27^Kip1^ expression (**Figure 7H**). Therefore, the interaction between the MB134-T cells and the substrate surface is a key determinant of ILC dormancy. The absence of this interaction, either physically or pharmacologically, resulted in a more pronounced dormant state. Clinically, these findings underscore the significance of low-adhesive microenvironments in inducing ILC dormancy, such as those experienced in circulation, prompting further investigations into ILC CTCs and therapeutics to restrict DTC interactions with the ECM.

### Microcuvette cultured cells yield distinct transcriptomic profiles

The transcriptomic profiles of MB134-P and MB134-T cells were screened across FN1 and COL3A1 micropatterns and microcuvettes under deprivation and control conditions. The transcriptomic profiles revealed stark differences, particularly between MB134-P and MB134-T cells (**Figure 8A**). Interestingly, the most differential profile was that of the cells on microcuvettes under deprivation conditions (**Figure 8A**). Under control conditions, the MB134-T cells had distinct expression profiles depending on the substrate, with 50 unique transcripts for FN1 micropatterns, 109 unique transcripts for COL3A1 micropatterns, 750 unique transcripts for the microcuvettes, and 52 shared transcripts between all three substrates (**Figure 8B**). Under deprivation conditions, the MB134-T cells exhibited a few more distinct transcripts, depending on the substrate, with 60 for FN1 micropatterns, 116 for COL3A1 micropatterns, 934 for microcuvettes, and 20 shared transcripts among the three (**Figure 8C**). Kyoto Encyclopedia of Genes and Genomes (KEGG) analyses were performed to determine pathways based on the unique and shared transcripts. For the shared transcripts under control conditions, several cell cycle regulatory genes were highlighted, including *CDK2*, *CCNB2*, and *PLK1* (**Figure 8D**), which were predominantly downregulated in deprived MB134-T cells. Interestingly, the pathways that were upregulated in microcuvettes under deprivation conditions were miRNA in cancer, polycomb repressive complex, and protein processing in the endoplasmic reticulum (**Figure 8E**). These results corroborate our previous findings that miRNA and histones targeted by the polycomb repressive complex contribute to the epigenetic induction of dormancy. On the other hand, endoplasmic reticulum stress is associated with dormancy in drug-resistant cancer cells,^78^ which is involved in protein processing within the endoplasmic reticulum. Lastly, 555 transcripts were specific to MB134-T cells compared to MB1314-P cells on the microcuvettes under deprivation (**Figure 8F**). The 555 transcripts were screened to identify those that coincided with the expression of p27^Kip1^ across the different substrates. A total of 13 transcripts correlated positively with the trends in p27^Kip1^ expression. A protein-protein interaction network analysis (STRING) was performed on these 13 transcripts and our previously identified differentially regulated proteins, revealing a network that included *MUC1* and *SAV1*, with *CDH1* at the center of the dormancy response (**Figure 8G**). Of the other 11 transcripts not included in the STRING analysis, some were implicated in cell cycle arrest (*UBXN2A*, *MXD4*, and *USP42*),^79-81^ endoplasmic reticulum stress response (*ERN1*),^82^ estrogen receptor alpha inhibition (*ZNF496*),^83^ actin cytoskeletal function and mechanosensing (*TPM2*),^84,85^ and early dissemination (*LTBP3*).^86^ Interestingly, *MUC1* is associated with tamoxifen resistance in breast cancer,^87^ while *SAV1* is implicated in cell cycle exit^88^ and is a key component of the Hippo pathway. These transcripts may reveal novel pathways by which ILC cells, particularly those resistant to anti-estrogen treatment, acquire a dormant phenotype.

**Figure 8:**
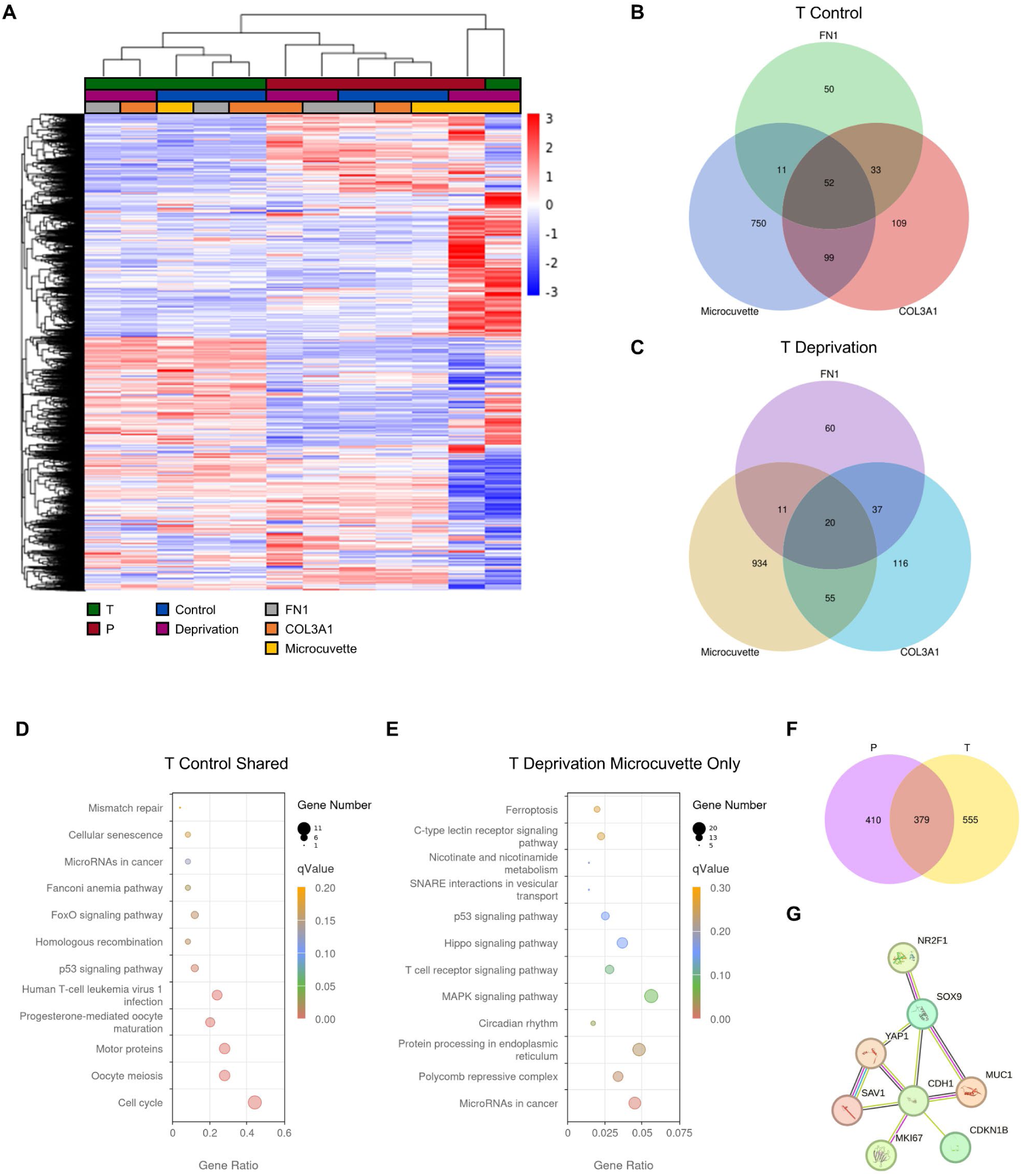
RNA sequencing across micropatterns and microcuvettes. (**A**) A heatmap exhibits the differences between groups that are color-coded (green represents MB134-T, red represents MB134-P, blue represents control conditions, magenta represents deprivation conditions, gray represents FN1 micropatterns, orange represents COL3A1 micropatterns, and yellow represents microcuvettes. (**B**) The Venn diagram illustrates unique or shared transcripts across FN1 micropatterns, COL3A1 micropatterns, and microcuvettes under control conditions. (**C**) The Venn diagram illustrates unique or shared transcripts across FN1 micropatterns, COL3A1 micropatterns, and microcuvettes under deprivation conditions. (**D**) KEGG analyses of the shared transcripts under control conditions. (**E**) KEGG analyses of the unique transcripts in microcuvettes under deprivation conditions. (**F**) The Venn diagram illustrates the unique or shared transcripts between MB134-P and MB134-T cells on the microcuvettes under deprivation conditions. (**G**) STRING analyses reveal a network of transcripts, including previously identified proteins and newly discovered transcripts derived from RNA sequencing. N = 2 for each condition.

## Discussion

Within this investigation, we employed multiple dimensions of cell culture through bioengineered approaches to examine various aspects of ILC biology and dormancy. Two-dimensional models enable the rapid, facile, and controllable actuation of cell-substrate interactions that are easily interpretable and maintainable, but lack *in vivo* relevance due to the homogeneity of nutrients, cell polarization, and limited cell-cell interactions.^89^ Three-dimensional models recapitulate the complexities of *in vivo* cell-cell and cell-substrate interactions, but are less tunable, reproducible, and manageable compared to two-dimensional assays.^89^ Pseudo-three-dimensional models, while not entirely three-dimensional due to some cell polarization, offer more ease than three-dimensional models and more *in vivo* relevance than two-dimensional models.^90^ Our two-dimensional protein micropatterns revealed profound differences in cell-substrate interactions between MB134-P and MB-134-T cells, which were more subtle in our three-dimensional and pseudo-three-dimensional methods. Moreover, the interaction with the two-dimensional surface influenced the dormancy response in MB134-T cells, exhibiting a dependence on the micropatterned protein, a mesenchymal-like phenotype, and filopodial formation when the cells were in a dormant state. Despite lacking physiological complexity, various bioengineered two-dimensional methods have equally advanced the field of breast cancer dormancy.^31^ To name a few, the covalent binding of ECM proteins to a rigid coverslip revealed the significance of fibronectin secretion and organization,^49^ tuning hyaluronic acid hydrogels with varying stiffnesses yielded a dormant phenotype on soft surfaces,^36^ and an organotypic microvascular niche co-culture demonstrated that endothelial cells induce dormancy.^91^ On the other hand, MB134 and MCF7 cells cultured in the three-dimensional, bioinert, agarose microgels resembled ductal and lobular morphology the most compared to the other culturing modalities. Although viability was compromised under deprivation conditions, the MB134-T cells were unable to form spheroids, whereas the MB134-P cells were able to form spheroids to some extent. Highly crosslinked three-dimensional hydrogels restricting cell adhesion induced apoptosis in breast cancer cells, which was rescued by the addition of integrin ligands, resulting in a viable dormant phenotype that was unable to form spheroids.^37^ In the absence of a confining matrix, cells in the pseudo-three-dimensional microcuvettes remained viable and induced a stronger dormancy phenotype in the MB134 cells compared to the two-dimensional protein micropatterns. Therefore, each dimension of culture imparts valuable insight with specific benefits and disadvantages.

Herein, we demonstrate that ILC dormancy is a multifaceted phenomenon centered around p27^Kip1^ signaling, with several potential perturbations that may facilitate the transition from dormancy to micrometastasis. The miRNAs, hsa-miR-221-3p and hsa-miR-222-3p, were downregulated in the dormancy-inducing condition for the MB134-T cells. Extracellular vesicles (EVs) are notorious for packaging miRNA and transferring cargo in an endocrine or paracrine fashion.^92,93^ Thus, supplementation of hsa-miR-221-3p and hsa-miR-222-3p via EVs may epigenetically inhibit the translation of p27^Kip1^ and promote re-awakening. On the other hand, MB134-T cells appear to be chemically and mechanically sensitive to the protein and stiffness to which they are bound. The ECM is a dynamic entity that is altered with age,^94^ inflammation,^95^ and obesity,^96^ eventually leading to stiff, fibrotic microenvironments. YAP is a major transcription factor for mechanotransduction via the Hippo signaling pathway^74^ that binds to the promoter of p27^Kip1^ and negatively affects its expression.^97^ We observed that MB134-T cells with nuclear YAP repressed the expression of p27^Kip1^. Both physical and pharmacological removal of the interaction with the rigid micropatterned surface resulted in a deeper state of quiescence, characterized by a reduction in Ki-67. Insults by external agents can alter the ECM composition and directly impact dormancy through the Hippo signaling pathway. Tobacco smoke-induced formation of neutrophil extracellular traps proteolyze the ECM, leading to YAP-mediated re-awakening of dormant DTCs in breast cancer.^98^ Similarly, chemotherapy disrupts the ECM, promoting YAP signaling and allowing p27^Kip1^-positive DTCs to cycle into micrometastasis in colon cancer.^99^ Therefore, various autocrine, paracrine, endocrine, and environmental perturbations exist that can destabilize p27^Kip1^ expression and enable micrometastasis, informing potential therapeutic interventions for ILC late recurrences.

Three main strategies are proposed for the clinical treatment of dormancy, including targeting dormant DTCs via inherent pathways,^100^ sensitizing dormant DTCs to chemotherapy,^101^ and perpetuating dormancy.^53^ Despite advances in preclinical investigations, translation to the clinic remains a bottleneck, with few current clinical trials, albeit with significant strides that are revolutionizing cancer treatment.^102,103^ A benefit of using micro-compartmentalized dormancy models, such as those illustrated in this investigation that mimic DTC dormancy, is the high-throughput testing of multiple therapeutic agents simultaneously and over short timeframes. On the other hand, the micro-compartmentalization of cells, particularly in well-defined arrays, enables the visualization of the same cell over extended periods under treatment. These benefits are non-trivial and expensive in animal models. Combining the benefits of micro-compartmentalized *in vitro* methods to identify candidates for subsequent *in vivo* testing will accelerate the development of novel treatments for ILC dormancy. Future investigations will utilize this system to assess dormancy, proliferation, and viability in ILC cells following the treatment with potential candidates that reduce the previously mentioned possible perturbations to p27^Kip1^ signaling, including treatments such as miRNA inhibitors/mimics, FAK/Pyk2 inhibitors, targeting novel genes and pathways from our RNA sequencing data, and agents currently in clinical trials, either solely or in combination with chemotherapies.

Transcriptomic and bioinformatic analyses revealed various interesting pathways that coincide with our findings and provide novel insights that require future validation. Interestingly, the MB134-T cells developed actin fibers compared to the MB134-P cells, and under deprivation conditions, formed actin-rich protrusions reminiscent of filopodia. *TPM2* stabilizes actin stress fibers^84^ and influences mechanosensing, suppressing growth on soft matrices.^85^ This corroborates our finding that MB134-T cells, when removed from a rigid adhesive substrate to a softer non-adhesive substrate, lose their proliferative capacity. Given that actomyosin contractility is central to ILC biology,^104,105^ the duality between extracellular sensing and sarcomere-like dynamics may provide unique insights into ILC dormancy. On the other hand, the STRING analysis illustrated *CDH1* as central to the ILC dormancy response, interconnected by mechanosensing, mesenchymal-like transition, and the cell cycle. Loss of *CDH1* is a significant mechanism for estrogen-receptor-positive breast cancer dormancy.^67^ With *CDH1* already deleted in ∼90% of patients with ILC^15^ and dormancy-tending cancer cells originating from the primary tumor,^48,52^ patients with ILC may be at a higher risk for developing dormant DTCs because of this deletion. These differences highlight the need to investigate ILC as separate from IDC, which infrequently harbors the *CDH1* deletion.

While the goal of this investigation was to circumvent the time-expensive experimentation of ILC murine models, a limitation of the study is the simplicity of our *in vitro* model, which lacks the complex heterogeneous interactions that occur in the pre-metastatic niche.^106^ Advances to the system include the utilization of organoids that harbor the complexities present *in vivo* through patient-derived samples,^107^ synthetic biology,^108^ or co-culturing with niche-resident cells.^109^ Furthermore, the cumulative stress of the microgel scaffold and the deprivation conditions resulted in cell death. Modifications to the microgel system to improve viability may include the addition of cell-binding sites, ECM proteins, or the aforementioned advancements, which could bring further complexity to our model.

Altogether, we demonstrated the ability of the bioengineered micro-compartmentalization techniques to recapitulate differences in ILC and IDC morphology, simulate phenotypes observed in dormant DTCs, and delineate differences in the ILC dormancy response. We found that the propensity to enter a dormant phenotype was influenced by tamoxifen-resistance-associated miRNA, with MB134-T cells demonstrating the highest potential for this phenotype. Furthermore, MB134-T cells acquired a stem-cell-like phenotype and were sensitive to both chemical and mechanical stimuli, which reflected their level of quiescence. We hope that this investigation will catalyze the field to further investigate ILC dormancy, an aspect of ILC progression that concerns a significant population of underrepresented patients who deserve greater attention in research.

## Materials and Methods

### Cell Culture

MCF7 cells (American Type Culture Collection; ATCC), HEK293T (ATCC) cells, and MDA-MB-231 cells (ATCC) were cultured in DMEM (Gibco) containing 10% (v/v) fetal bovine serum (FBS; Gibco) and 1% (v/v) penicillin-streptomycin (Gibco). MDA-MB-134 VI cells (ATCC) were cultured in a 1:1 mixture of DMEM:L15 (Gibco) containing 10% (v/v) FBS and 1% (v/v) penicillin–streptomycin. SUM44PE cells (Asterand) were grown Ham’s F12 media (Gibco) supplemented with 1 mg/mL of bovine serum albumin (BSA; Sigma-Aldrich), 5 mM of ethanolamine (Sigma-Aldrich), 10 mM of a N-2-hydroxyethylpiperazine-N’-2-ethanesulfonic acid (HEPES) buffer (Sigma-Aldrich), 1 µg/mL of hydrocortisone (Sigma-Aldrich), 5 µg/mL of insulin (Sigma-Aldrich), 50 nM of sodium selenite (Sigma-Aldrich), 5 µg/mL of apo-transferrin (Sigma-Aldrich), and 10 nM of triiodo thyronine (Sigma-Aldrich). The cells were maintained in a 5% CO_2_ incubator at 37 °C and 21% O_2_ (Thermo Scientific). The cells were detached with TrypLE™ Express Enzyme (Gibco) for subculturing and before experimentation. Tamoxifen-resistant cells were generated by culturing the MDA-MB-134 VI cells in the presence of 100 nM of 4-hydroxy Tamoxifen (4-HT; Cayman Chemical) and SUM44PE cells in the presence of 500 nM of 4-HT for 6 months.^43^

### PIP-FUCCI Biosensor Infection

pLenti-PGK-Neo-PIP-FUCCI was a gift from Dr. Jean Cook (Addgene, plasmid #118616).^51^ Competent *E. coli* cells (DH5α; Invitrogen) were transformed using the plasmid and single colonies of the transformed bacteria were selected from Luria Broth (LB)-agar plates (Fisher Scientific) for further expansion in bacterial broth, containing 1% (w/v) tryptone (Fisher Scientific), 1% (w/v) NaCl (Fisher Scientific), 0.5% (w/v) yeast extract (Fisher Scientific), and 50 µg/mL of ampicillin (Sigma-Aldrich). DNA was extracted with the QIAprep Spin Miniprep Kit (Qiagen), according to the manufacturer’s instructions. The plasmid was confirmed using restriction enzyme digestion. For virus production, HEK293T cells were transfected using P3000™ Reagent (Invitrogen) with a mixture of pLenti-PGK-Neo-PIP-FUCCI, PAX2, and PMD2 plasmids at a 2:1:1 ratio in OptiMEM™ media (Invitrogen), following the manufacturer’s protocol. After 48 hr of incubation, the viral supernatant from the HEK293T cells was used to infect MB134-P and MB134-T cells with 8 µg/ml of polybrene (Sigma-Aldrich). The infection was repeated twice, after which the cells were expanded and sorted with flow cytometry.

### Light-Induced Molecular Adsorption (LIMA)

Glass coverslips (Globe Scientific) were cleaned in 70% ethanol (Decon Labs) and DI water for 3 min each in a sonication bath. The clean coverslips were subsequently treated with oxygen plasma (Plasma Etch) to clean the surface further and induce a negative charge. The plasma-treated coverslips adhered reversibly to silicon gaskets with 7 mm × 7 mm wells (Grace BioLabs). 50 μL of 0.01% (w/v) PLL (Sigma-Aldrich) was adsorbed onto the surface for 1 hr at room temperature, followed by three washes with 100 μL of 0.1 M HEPES buffer (pH = 8.5; Boston BioProducts). A solution of 100 mg/mL of mPEG-SVA (Laysan Bio) in HEPES buffer was quickly added to the PLL-coated coverslip and allowed to react for 1 hr at room temperature. The PLL-*g*-mPEG-coated surface was washed with DI water, followed by a coating with a benzophenone-based photoinitiator gel (Alvéole). The functionalized surface was then exposed to 30 mJ/mm^2^ of UV light, following a template of 100-µm diameter circles spaced 200 µm center-to-center using a DMD-based projection system (Alvéole). The masks were generated via an in-house MATLAB (MathWorks) microarray generator. Following the photochemical reaction, the remaining photoinitiator was washed with DI water. The photopatterned surface was rehydrated in phosphate-buffered saline (PBS; Gibco) or a solution of 0.01 N HCl (Fisher Chemical) for 5 min, depending on the choice of ECM protein. For FN1 (Corning) adsorption, a 50 µg/mL solution in PBS was added to the photopatterned surface for 5 min. For COL3A1 (Advanced BioMatrix) adsorption, a 50 µg/mL solution in 0.01 N HCl was added to the photopatterned surface for 5 min. The ECM micropatterns were washed three times with PBS and twice with media. Cells were added at a concentration of 1 × 10^6^ cells/mL for 15 min and then washed with PBS to remove unadhered cells.

### *In Situ* Microcuvette Generation

A circular well of PDMS (The Dow Chemical Company) with a depth of 200 µm was placed on glass-bottom 24-well plates (Cellvis) on top of which another 200-µm PDMS ceiling was placed that contained an inlet and an outlet. The sealed circular well was designed to minimize the formation of bubbles. A 5% (w/v) solution of 4-arm PEGA (Laysan Bio) in 14.5 mg/mL of the photoinitiator (Alvéole) was added to the well and exposed to 120 mJ/mm^2^ of UV light via the DMD-based projection system (Alvéole). The mask was designed as a parabolic gradient with a diameter of 150 µm spaced 150 µm center-to-center. Following UV illumination, the microcuvettes were washed with 70% (v/v) ethanol, followed by PBS, and then with the media. The cells were added to the microcuvettes at a concentration of 1 × 10^6^ cells/mL.

### Microfluidic Generation of Cell-Laden Microgels

Previously reported cell encapsulation devices^41^ were coated with Aquapel (Pittsburgh Glass Works) to provide hydrophobicity to the microchannels. For the dispersed phase, ultra-low gelling temperature agarose (Sigma-Aldrich) was dissolved in PBS at a concentration of 2% (w/v) via microwave-induced heating, and then cooled to 37 °C. Half of the agarose solution was diluted to 1% (w/v) in PBS. The other half was diluted to 1% (w/v) in a cell suspension of 1 × 10^7^ cells/mL for long-term experiments or 3 × 10^7^ cells/mL for short-term experiments (to match the number of cells in both micropattern- and microcuvette-based micropatterning). A 1% (v/v) Pico-Surf™ 1 (Sphere Fluidics) solution in Novec™ 7500 (3M) was prepared for the continuous phase. The device was wet with the continuous phase via the top inlet, followed by the introduction of the dispersed phase, containing 1% (w/v) agarose, through the middle inlet, and the dispersed phase containing cells through the bottom inlet. The microdroplets were collected for 10 min in an ice bath, after which the emulsification was further allowed to crosslink for 20 min at 4 °C. Following gelation, the remaining Novec™ 7500 was removed. A 20% (v/v) solution of PFO (Sigma-Aldrich) in Novec™ 7500 was added to the cell-laden microgels to facilitate the exchange to an aqueous buffer. The PFO solution was removed, and the cell-laden microgels were resuspended in PBS. The colloid was centrifuged at 800 rpm for 30 seconds to remove all other non-polar impurities. The cell-laden microgels were washed three times with PBS on a 70 μm filter and transferred to the media.

### Inducing Dormancy

The cells undergoing growth factor and oxygen deprivation were incubated in 1% O_2_, 5% CO_2_, and 37 °C in serum-free media. On the other hand, the control was incubated in 21% O_2_, 5% CO_2_, and 37 °C in 10% (v/v) FBS. For IFN-γ-induced dormancy, 100 ng/mL of recombinant human IFN-γ (Peprotech)^41,110^ was added to the media, and the cells were incubated at 21% O_2_, 5% CO_2_, and 37 °C in 10% (v/v) FBS. The control was incubated in 21% O_2_, 5% CO_2_, and 37 °C in 10% (v/v) FBS without IFN-γ.

### Cellular Viability

The viability of the cells was assessed following deprivation treatment using the LIVE/DEAD® Viability/Cytotoxicity Kit (Invitrogen). The cells were incubated in PBS containing 2 μM of calcein AM and 4 μM of ethidium homodimer-1 for 30 min before imaging.

### Quantitative Reverse-Transcription Polymerase Chain Reaction (qRT-PCR)

According to the manufacturer’s instructions, RNA was extracted from the cells via the RNeasy Mini Kit (Qiagen). cDNA and amplification were performed using the TaqMan MicroRNA Assay (Applied Biosystems), following the manufacturer’s protocol. qRT-PCR (Applied Biosystems) was conducted for 40 cycles and normalized to GAPDH. The primers utilized for qRT-PCR are provided in **Supplementary Table 1**.

### Immunofluorescence

The samples were incubated in PBS containing 4% (v/v) paraformaldehyde (Cell Signaling Technology) for 15 minutes at room temperature. Following fixation, the samples were washed three times with PBS for 5 min each. All washing steps were performed as such. The samples were then permeabilized with 0.1% (v/v) Triton^TM^ X-100 (SAFC) in PBS for 10 min at room temperature, followed by washing with PBS. The samples were blocked for 1 hr in 10% (w/v) normal goat serum (NGS; Invitrogen) at room temperature, followed by incubation in the primary antibody (**Supplementary Table 2**) diluted in a 1% (w/v) BSA solution in 0.07% (v/v) Tween® 20 (Sigma-Aldrich) in PBS (PBS-T) and incubated overnight at 4 °C. The next day, the samples were then washed twice with PBS and once with PBS-T. The samples were then incubated with the secondary antibody diluted in a 1% (w/v) BSA solution in PBS-T for 1 hr at room temperature, followed by three PBS washes at 5-min intervals. The samples were then incubated with rabbit IgG (11.4 μg/mL; Invitrogen) in 10% (v/v) NGS for 1 hr at room temperature to block all secondary sites. The samples were then incubated with a primary fluorophore-conjugated antibody diluted in a 1% (w/v) BSA solution in PBS-T overnight at 4 °C. For actin staining, the ActinRed^TM^ 555 ReadyProbes^TM^ (Invitrogen) reagent was used according to the manufacturer’s protocol. All cells were then counterstained with 300 nM 4’,6-diamidino-2-phenylindole (DAPI; Invitrogen) for 5 min, after which the solution was washed away with PBS. For LIMA, buffer exchanges were performed via quick aspiration and application. ProLong^TM^ Glass Antifade Mountant (Invitrogen) was used next to mount the glass slides with stained cells. For the microgels, buffer exchanges were performed via centrifugation at 800 rpm for 5 min. The colloid was immobilized in a 1% (w/v) agarose solution in PBS by allowing the microgels to settle to the bottom of the coverslip at 37 °C. Afterward, the colloid solution was gelled at 4 °C for 20 min.

### Microscopy

Various microscopy methods were utilized in the study. For live-cell imaging and most quantitative purposes, a combination of epifluorescence and brightfield illumination was used, utilizing 10×, 20×, and 40× objectives in the Nikon Eclipse Ti2 microscope (Nikon Instruments). For live-cell imaging, the surrounding environment was maintained at 37 °C and 5% CO_2_ with 21% O_2_ for normoxic conditions and 1% O_2_ for hypoxic conditions, while utilizing laser powers below 10% and exposure times below 200 ms. For three-dimensional reconstruction, an Olympus FV3000 confocal microscope (Olympus) was used at 60 × oil-immersion magnification, capturing *z*-stacks that were compiled to generate three-dimensional images. On the other hand, microcuvette three-dimensional reconstruction was further performed using a Nikon® AXR confocal microscope (Nikon Instruments) with a piezo drive at 20 × magnification, during which *z*-stacks were captured and compiled to generate three-dimensional images. To visualize the coverslip surface, a Nikon® Ti2 microscope was used for TIRFM with an oil-immersion 100× objective. Background fluorescence was removed via rolling ball image subtraction. For scanning electron microscopy (SEM), a stepwise ethanol dehydration process was implemented, involving sequential immersions in 75%, 85%, 90%, and 100% ethanol for 10 min each. Subsequently, the coverslips were swiftly transferred to hexamethyldisilazane (HMDS; Sigma-Aldrich) for 10 min. Following the HMDS treatment, the coverslips were air-dried overnight before imaging. SEM images were taken using an Apreo SEM (Thermo Scientific) at 5.00 kV and a 25 pA current at a working distance of 9.9 mm.

### Image Analysis

Images were quantified utilizing various manual and automatic approaches with Fiji (ImageJ), NIS-Elements imaging software (Nikon Instruments), and MATLAB. For percent positivity or the total number of cells, positive cells were determined by equalizing the look-up tables (LUTs) and manually counting the high-expressing cells for each micropattern. For less pronounced binary expressions, LUTs were equalized, regions of interest (ROI) were generated that encompassed nuclear signals as determined by DAPI staining, and the mean fluorescent signal for each micropattern was measured. For measuring the characteristics of micropatterns or cells, thresholding was used to generate ROIs, the shape descriptors of which were then measured. In-house MATLAB algorithms were also utilized for live-cell tracking. Spot tracking on the NIS-Elements software was used to track changes in the biosensor-expressing cells.

### RNA Sequencing

Total RNA was extracted from cultured cells using the RNeasy Plus Mini Kit (Qiagen), according to the manufacturer’s instructions. The media was removed from the bioengineered models and transferred with 350 µL of Buffer RLT Plus containing 1% β-mercaptoethanol by vigorous pipetting until cells were released entirely. The lysate was transferred to a gDNA Eliminator spin column and centrifuged at 8000 × *g* for 30 seconds to remove genomic DNA. The flow-through was mixed with an equal volume of 70% ethanol, gently vortexed, and applied to an RNeasy spin column. After centrifugation at 8000 × *g* for 15 seconds, the flow-through was discarded. The column-bound RNA was washed by adding 700 µL of Buffer RW1, followed by 500 µL of Buffer RPE, with centrifugation between each step. A final centrifugation at full speed for 1 minute was performed to dry the membrane. RNA was eluted in 50 µL of RNase-free water by centrifugation at 8000 × *g* for 1 minute. RNA concentration and purity were assessed using a NanoDrop spectrophotometer. RNA samples were processed by Novogene for RNA sequencing and analyzed by NovoMagic.

### Statistical Analyses

Analysis of variance (ANOVA) was utilized for multiple group comparisons, with pairwise comparisons performed within the procedure. Two-tailed or paired *t*-tests were performed for two-sample comparisons. Holm’s procedure was employed to correct for multiple comparisons. All data are represented as the mean ± the standard error of the mean. Statistical comparisons are illustrated with asterisks or other symbols, with standalone asterisks denoting significance against their respective control and other symbols denoting significance between groups, not individual comparisons, unless stated otherwise.

## Acknowledgment

We thank Anthony McCoy, Michael Wilson, and Leigh Evrard for their assistance. Electron microscopy was performed at the Center for Electron Microscopy and Analysis (CEMAS) at The Ohio State University. Schematics were created with BioRender.com. We acknowledge resources from the Campus Microscopy and Imaging Facility (CMIF) at The Ohio State University.

## Author contributions

Conceptualization: ER, BR, XYR, SM. Methodology: ER, XYR. Software: XYR. Investigation, data curation, and validation: XYR, SM, CH, DSP, HL, XH, KTN, CKN, MHD. Formal analysis: XYR, SM, JDR, XZ. Resources and funding acquisition: ER, BR, XZ, DGS, SM, SMM, XYR. Writing – original draft preparation: XYR. Writing – review and editing: ER, BR, DGS, SM, XZ, ES, SMM, XYR. Visualization: XYR, ER, SM, BR. Supervision/ Project administration: ER, BR.

## Data and materials availability

All data needed to evaluate the conclusions in the paper are present in the paper and/or the Supplementary Materials.

