## Supplementary Materials for "Multidimensional Cellular Micro-Compartments to Model Invasive Lobular Carcinoma Dormancy"

**Supplementary Table 1: List of qRT-PCR primers**

| Target | Vendor | Assay ID |
| --- | --- | --- |
| hsa-miR-221-3p | Thermo Fisher Scientific | 000524 |
| hsa-miR-222-3p | Thermo Fisher Scientific | 002276 |
| GAPDH | Thermo Fisher Scientific | Hs02786624_g1 |

**Supplementary Table 2: List of antibodies**

| Target | Vendor | Catalog Number | Dilution |
|---|---|---|---|
| Ki-67 | Cell Signaling Technology | 9129S | 1:400 |
| $p21^{Cip1/Waf1}$ | Cell Signaling Technology | 5487S | 1:400 |
| $p27^{Kip1}$ | BD Biosciences | 610241 | 1:500 |
| NR2F1 (COUP-TF1) | Abcam | ab211712 | 1:500 |
| H3K27me3 | Cell Signaling Technology | 9733S | 1:500 |
| E-Cadherin | Cell Signaling Technology | 3195S | 1:100 |
| $p16^{Ink4}$ | Cell Signaling Technology | 18769S | 1:400 |
| β-Tubulin | Cell Signaling Technology | 2128S | 1:200 |
| SOX9 | Millipore | AB5535 | 1:500 |
| YAP | Cell Signaling Technology | 38707S | 1:100 |

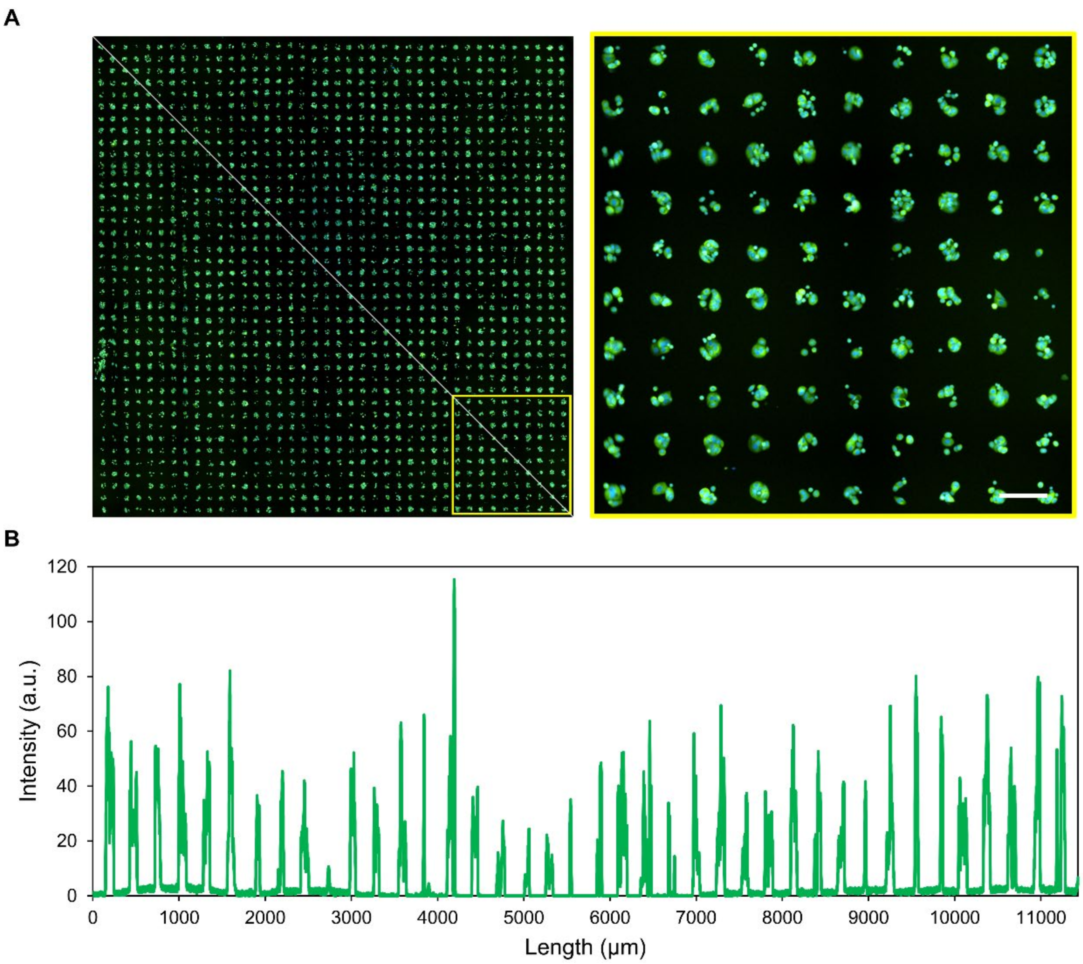


**Supplementary Figure 1: Large-scale micropatterning of cells on ECM proteins.**

(**A**) An epifluorescence image via automatic montage generation illustrates the reproducibility of the two-dimensional micropatterns. The yellow box is magnified in the right panel, revealing a 10 × 10 array of micropatterns. (**B**) The diagonal line is quantified for fluorescence as a function of length, demonstrating equally spaced patterns that are homogeneously filled. The scale bar is 200 μm.

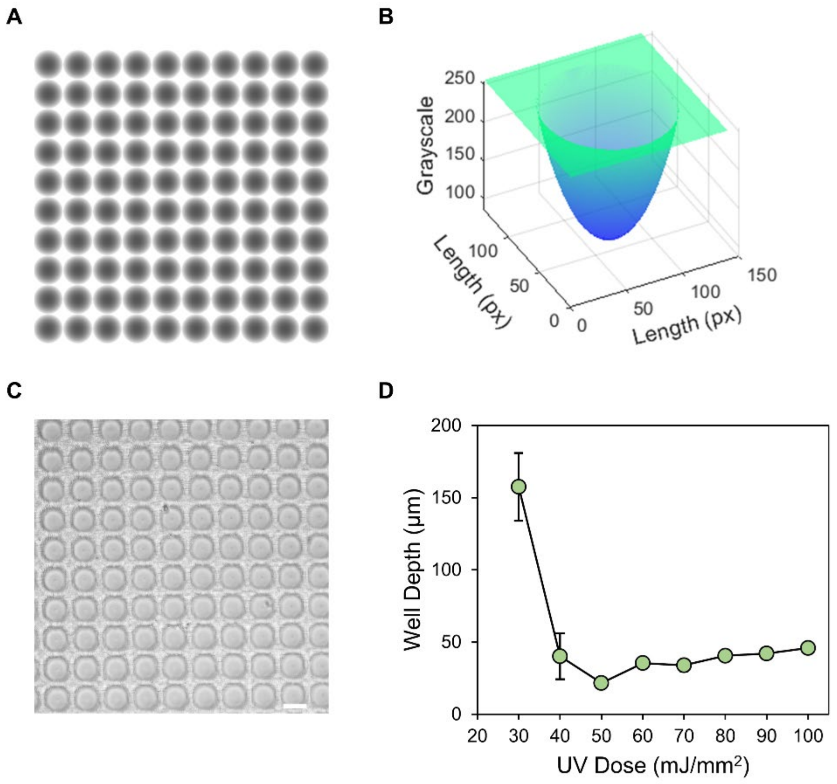


**Supplementary Figure 2: Large-scale elaboration of microcuvettes.**

(**A**) A grayscale digital image with a paraboloid array is rapidly converted to photon flux corresponding to the grayscale values with a DMD. (**B**) A surface plot of a microcuvette demonstrates that the grayscale follows a paraboloid as a function of the area. (**C**) Brightfield images of the microcuvettes illustrate their reproducibility. (**D**) The depth of the microcuvettes as a function of UV dose exhibits a plateau beyond 60 $mJ/mm^2$ (N = 3, error bars indicate the standard error of the mean). The scale bar is 100 µm.

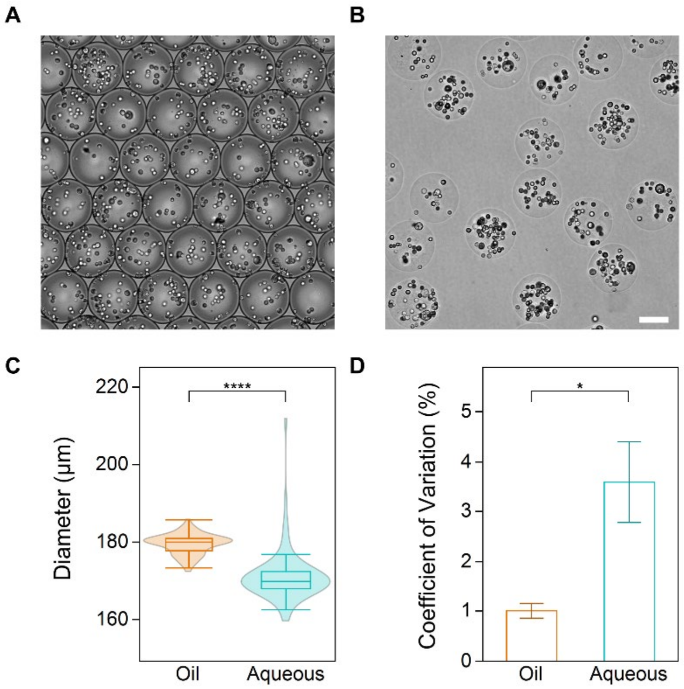


**Supplementary Figure 3: Large-scale generation of microgels.**

(**A**) Representative brightfield images of microdroplets in the oil phase demonstrate homogeneity. (**B**) Representative brightfield images of microgels in aqueous phase exhibit continued homogeneity despite phase transfer. (**C**) The diameter of the microdroplets in the oil and aqueous phase is represented as violin-box conjugate plots and demonstrates a reduction in diameter (N = 3, n = 25; ****$p < 0.0001$). (**D**) The coefficient of variation increases after transferring to the aqueous phase (N = 3, error bars indicate the standard error of the mean; *$p < 0.05$). The scale bar is 100 µm.

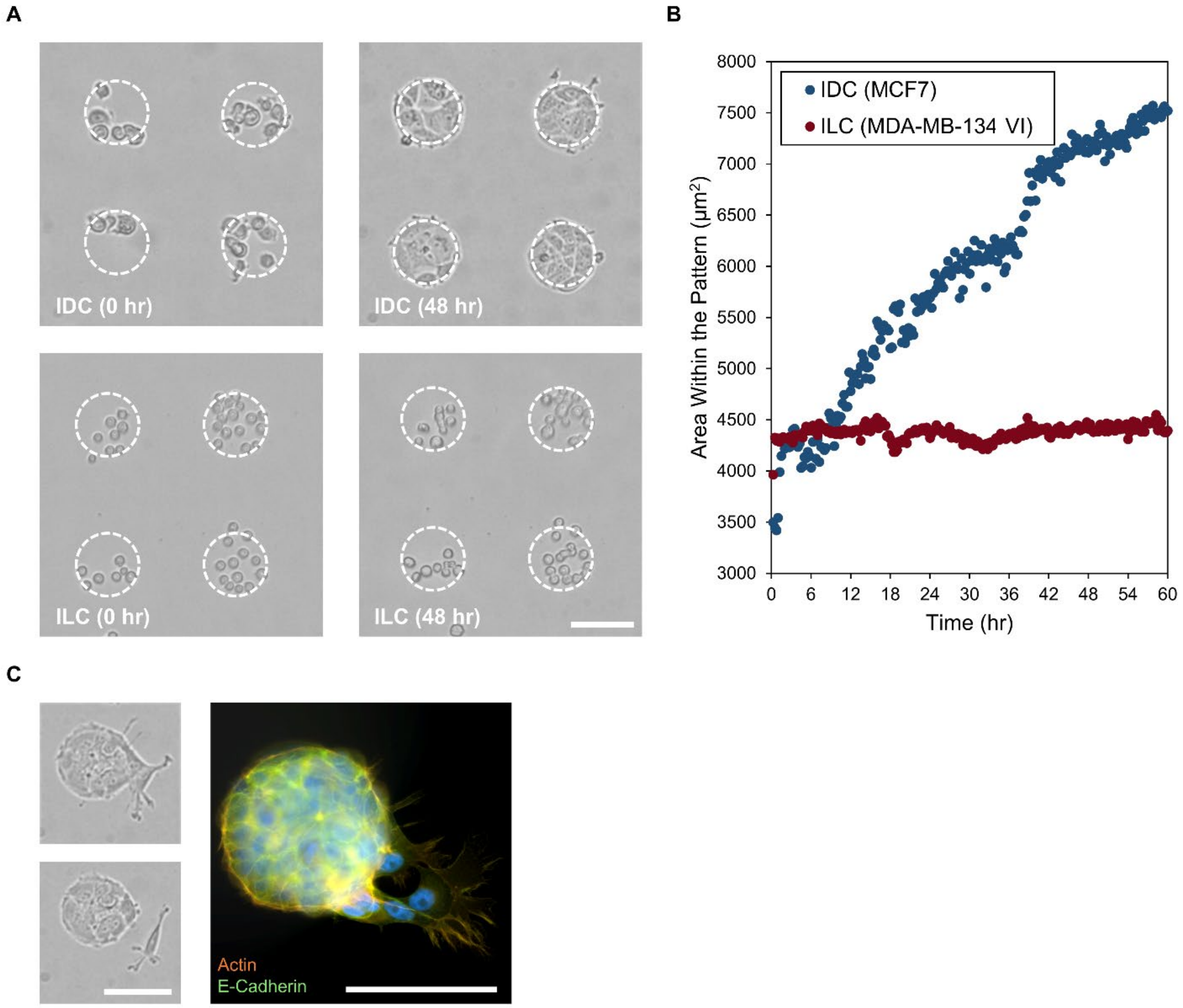


**Supplementary Figure 4: Cellular filling of FN1 micropatterns.**

(**A**) FN1 micropatterns seeded with a model IDC cell line (MCF7) and ILC cell line (MB134) exhibit differences in micropattern filling. (**B**) The average area within the pattern as a function of time demonstrates the differential filling of the MCF7 and MB134 cells (n = 9). (**C**) The MCF7 cells expel single cells as visualized with live-cell brightfield (left) and epifluorescence (right; green represents E-cadherin, orange represents actin, and blue represents DAPI). All scale bars are 100 μm.

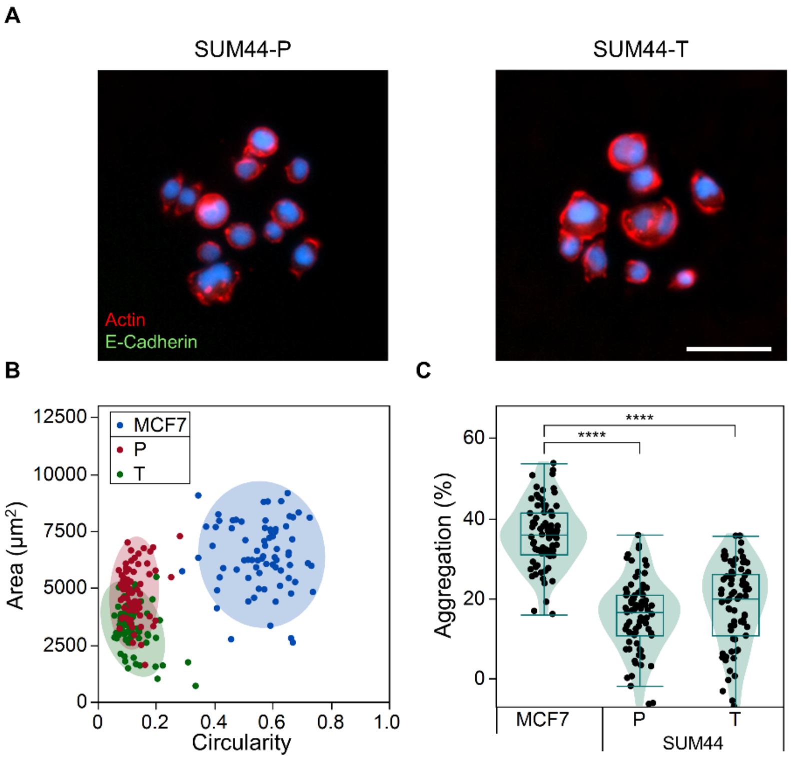


**Supplementary Figure 5: Similarities between SUM44-P and SUM44-T cells.**

(**A**) Epifluorescence microscopy of the cells on FN1 microdomains reveals differences in interactions with the micropatterned proteins and E-cadherin expression (green represents E-cadherin, red represents actin, and blue represents DAPI). (**B**) The interaction is depicted as clusters of the area *versus* the circularity of cellular coverage over the microdomain (N = 3, n = 25). (**C**) The aggregation of cells, depicted as violin-box conjugate plots, demonstrates differential aggregation (N = 3, n = 25; ****$p < 0.0001$). The scale bar is 50 μm.

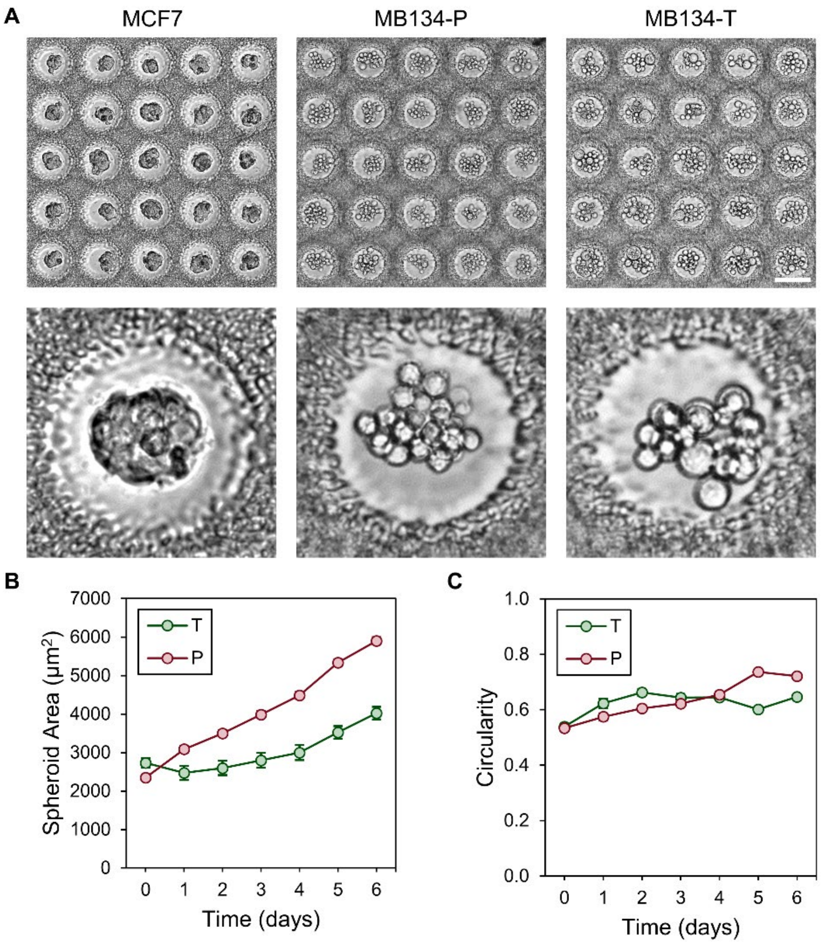


**Supplementary Figure 6: Long-term microcuvette culture.**

(**A**) Representative brightfield images of the MCF7, MB134-P, and MB134-T cells on the microcuvettes show differential aggregation after 12 hours. The bottom images show a single microcuvette. (**B**) The MB134-P and MB134-T spheroidal areas over 6 days reveal differences in growth (N = 3, n = 25, error bars indicate the standard error of the mean). (**C**) The MB134-P and MB134-T spheroidal circularity over 6 days reveals moderate to no increases in circularity (N = 3, n = 25, error bars indicate the standard error of the mean). The scale bar is 100 μm.

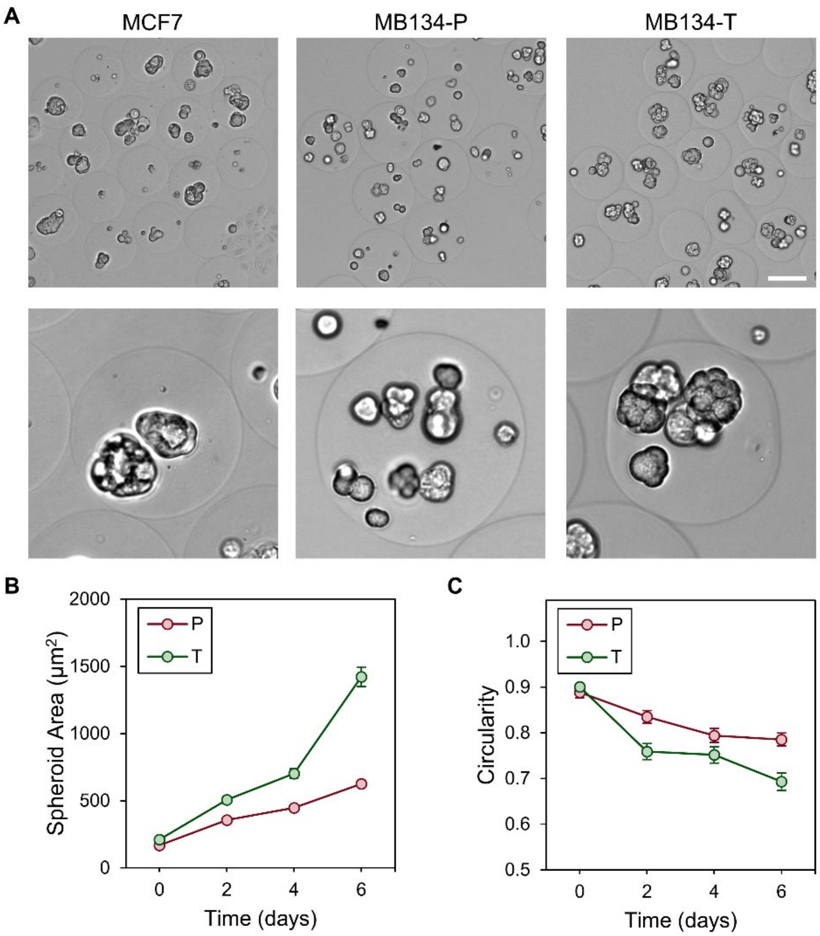


**Supplementary Figure 7: Long-term microgel culture.**

(**A**) Representative brightfield images of the MCF7, MB134-P, and MB134-T cells within microgels show differential morphologies after 6 days. The bottom images show a single microgel. (**B**) The MB134-P and MB134-T spheroidal areas over 6 days reveal differences in growth (N = 3, n = 25, error bars indicate the standard error of the mean). (**C**) The MB134-P and MB134-T spheroidal circularity over 6 days reveals decreases in circularity (N = 3, n = 25, error bars indicate the standard error of the mean). The scale bar is 100 μm.

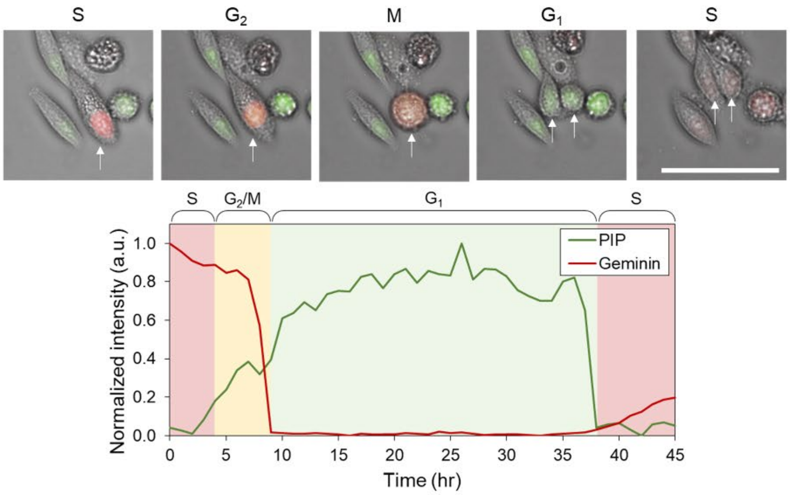


**Supplementary Figure 8: PIP-FUCCI to track the ILC cell cycle live.**

The biosensor was designed so that cells in the $G_1$-phase are visualized as green via the proliferating cell nuclear antigen (PCNA)-interacting protein (PIP) degron from Cdt1 ($Cdt1_{1-17}$), cells in the S-phase are visualized as red via $Geminin_{1-110}$, cells in the $G_2$-phase are visualized as yellow due to colocalization of both fluorescent proteins, and cells in M phase are visualized as yellow dispersed throughout the cell. The scale bar is 100 µm.

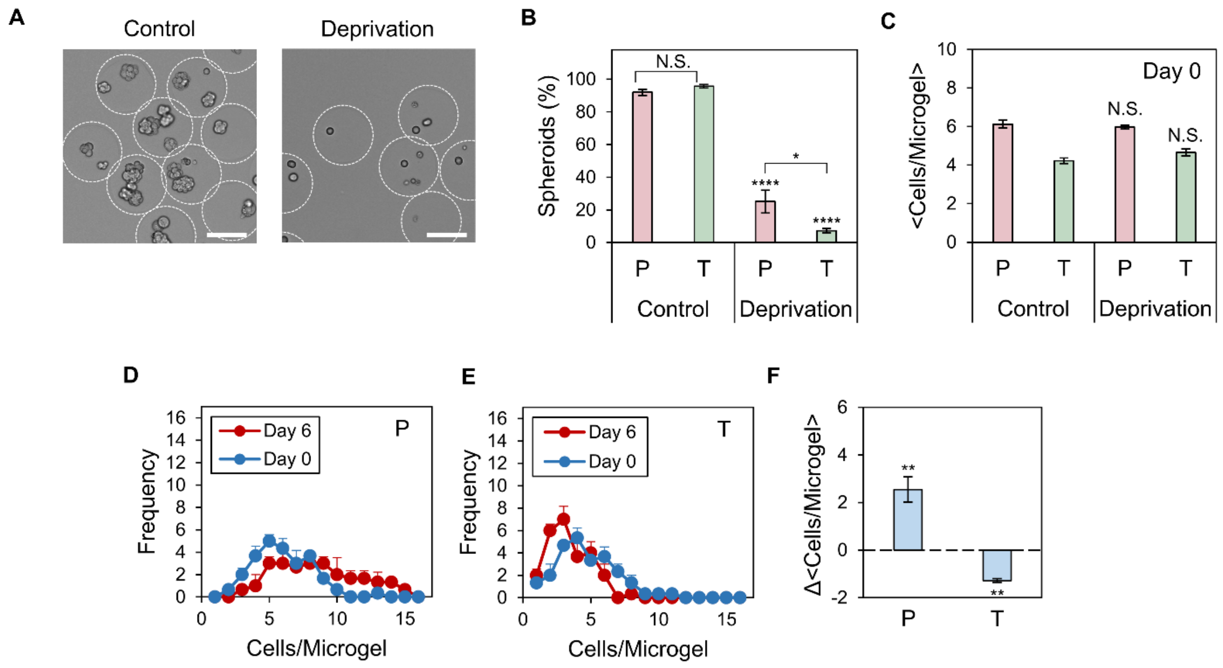


**Supplementary Figure 9: Microgels recapitulate single-cell dormancy despite low viability.** (**A**) Representative brightfield images of MB134-T cells under control and deprivation conditions illustrate the formation of spheroids in the former and single cells dispersed in the matrix for the latter. (**B**) The percentage of spheroids, represented as bar plots, demonstrates a loss of spheroids under deprivation conditions, which is more pronounced for the MB134-T cells (N = 3, error bars indicate the standard error of the mean; ****$p < 0.0001$, *$p < 0.05$). (**C**) Despite these observed differences, the cellular occupancy post-encapsulation was the same at day 0, indicating the differences are due to the deprivation conditions (N = 3, error bars indicate the standard error of the mean). (**D**) Histograms of cellular occupancy for the deprivation conditions at days 0 and 6 for the MB134-P cells show an increase in cell number (N = 3, error bars indicate the standard error of the mean). (**E**) Histograms of cellular occupancy under deprivation conditions at days 0 and 6 for the MB134-T cells demonstrate a loss of cells (N = 3, error bars indicate the standard error of the mean). (**F**) The change in the average cellular occupancy under deprivation conditions reveals an increase in MB134-P cells and a decrease in MB134-T cells (N = 3, error bars indicate the standard error of the mean; $^{**}p < 0.01$). All scale bars are 100 μm.

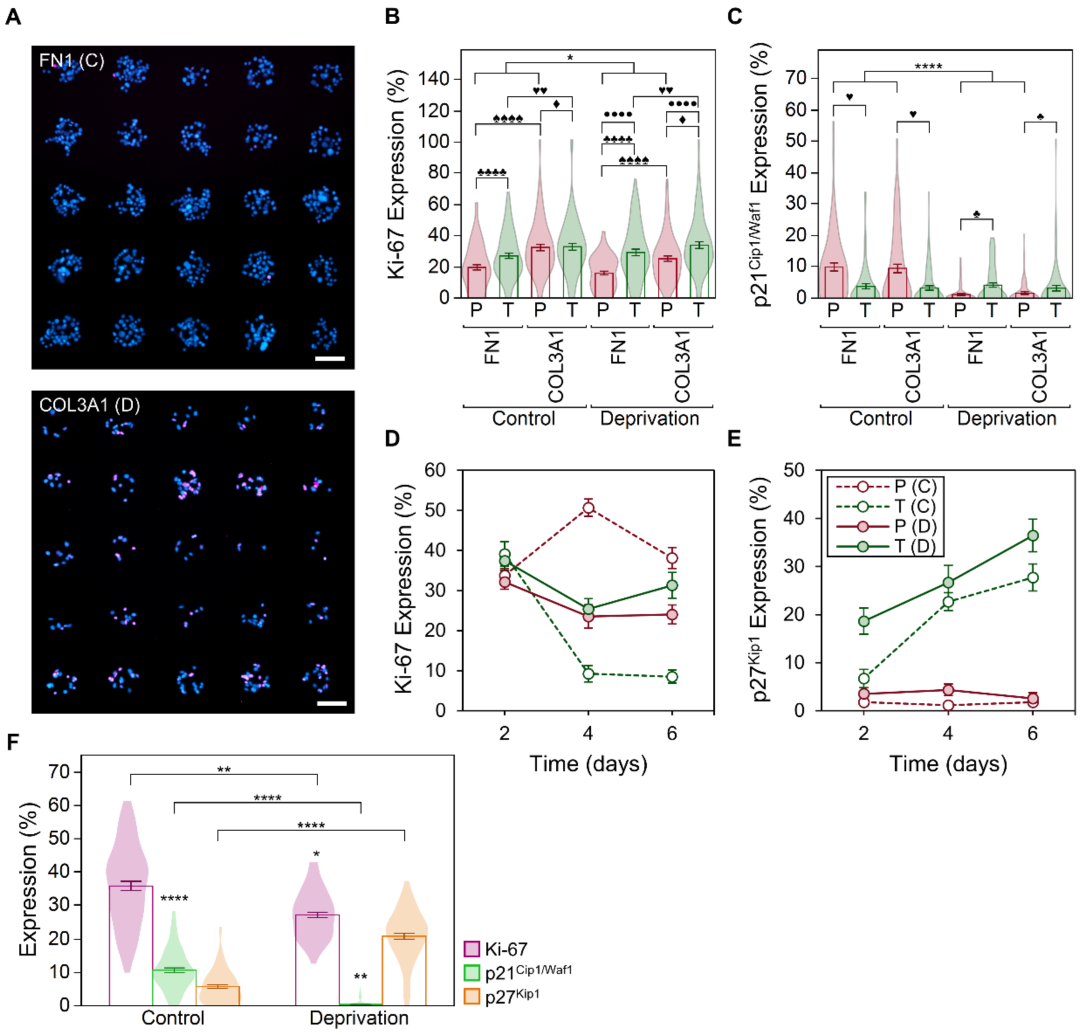


**Supplementary Figure 10: Effects of micropatterns and deprivation on MB134 cells.**

(**A**) Representative epifluorescence images of p27$^{Kip1}$ expression in an array illustrate the least-expressing (FN1 micropatterns, control) and most-expressing (COL3A1 micropatterns, deprivation) conditions (magenta represents p27$^{Kip1}$ and blue represents DAPI). (**B**) The expression of Ki-67 is depicted as violin-bar conjugate plots (N = 3, n = 25, error bars indicate the standard error of the mean; **p* < 0.05, ♦*p* < 0.05 for the group, ♥♥*p* < 0.01 for the group, ♣♣♣♣*p* < 0.0001 for the group, ♠♠♠♠*p* < 0.0001 for the group, ••••*p* < 0.0001 for the group). (**C**) The expression of p21$^{Cip1/Waf1}$ is depicted as violin-bar conjugate plots (N = 3, n = 25, error bars indicate the standard error of the mean; **p* < 0.0001, ♥*p* < 0.05 for the group, ♣*p* < 0.05 for the group). (**D**)

The long-term expression of Ki-67 in MB134-P and MB134-T cells under control and deprivation conditions over 6 days demonstrates that 2 days is the optimal condition for investigating dormancy (n = 25, error bars indicate the standard error of the mean). (**E**) The long-term expression of $p27^{Kip1}$ in MB134-P and MB134-T cells under control and deprivation conditions over 6 days demonstrates that 2 days is the optimal condition for investigating dormancy (n = 25, error bars indicate the standard error of the mean). (**F**) The expression profiles of Ki-67, $p21^{Cip1/Waf1}$, and $p27^{Kip1}$ as violin-bar conjugate plots on 200-μm patterns reveal similarities with 100-μm patterns (N = 3, n = 25, error bars indicate the standard error of the mean; $*p < 0.05$, $**p < 0.01$, $****p < 0.0001$, with standalone asterisks denoting a comparison to 100-μm patterns). All scale bars are 100 μm.

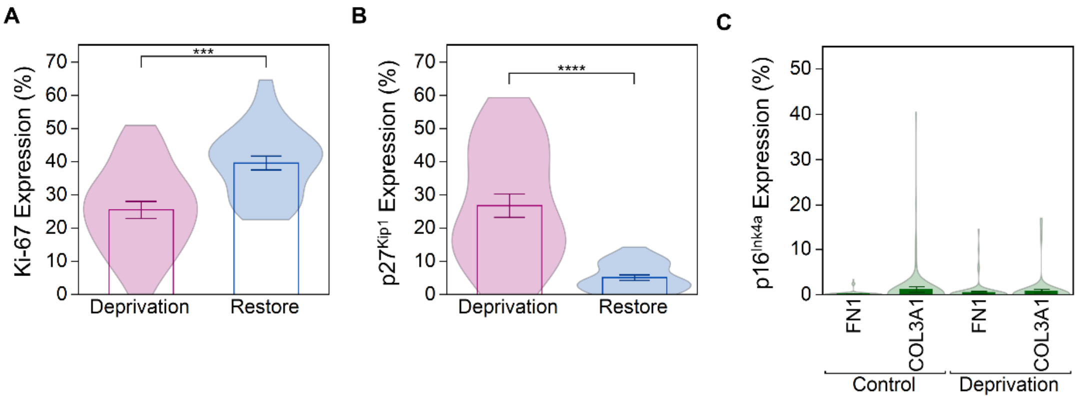


**Supplementary Figure 11: Reawakening and senescence in MB134-T cells.**

(**A**) The expression of Ki-67, depicted as violin-bar conjugate plots, is enhanced when control conditions are restored for 48 hr after MB134-T cells were in deprivation conditions for 48 hr, with the control being deprivation conditions for 96 hr (n = 25, error bars indicate the standard error of the mean; ***$p < 0.001$). (**B**) The expression of $p27^{Kip1}$, depicted as violin-bar conjugate plots, is reduced when control conditions are restored for 48 hr after MB134-T cells were in deprivation conditions for 48 hr, with the control being deprivation conditions for 96 hr (n = 25, error bars indicate the standard error of the mean; ****$p < 0.0001$). (**C**) The expression of $p16^{Ink4a}$ under standard deprivation conditions for 48 hr is depicted as violin-bar conjugate plots, revealing minimal expression across all conditions (N = 3, n = 25, error bars indicate the standard error of the mean).

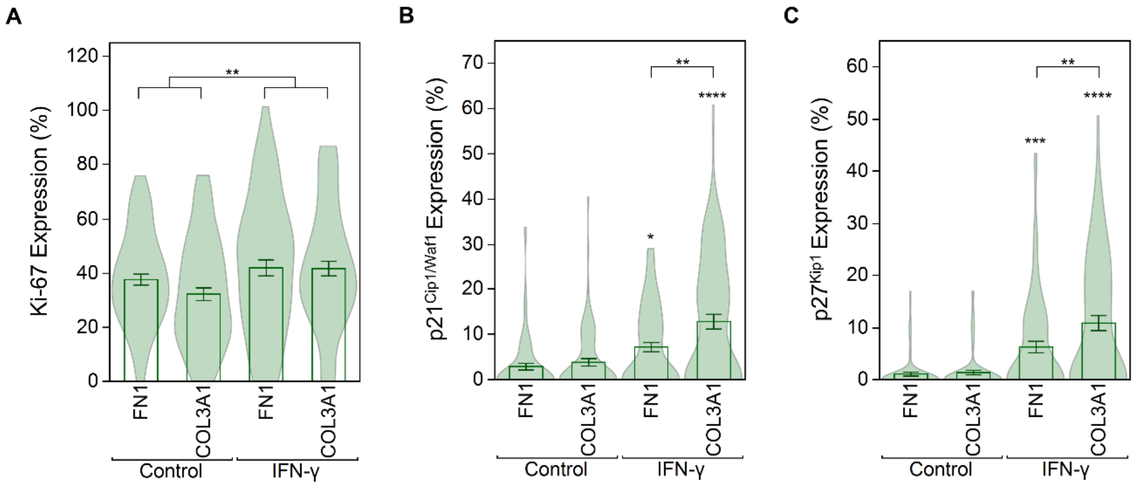


**Supplementary Figure 12: Effects of micropatterns and IFN-γ treatment on MB134-T cells.** (**A**) The expression of Ki-67 is depicted as violin-bar conjugate plots (N = 3, n = 25, error bars indicate the standard error of the mean; $^{**}p < 0.01$). (**B**) The expression of $p21^{Cip/Waf1}$ is depicted as violin-bar conjugate plots, revealing increased expression with COL3A1 micropatterns under treatment (N = 3, n = 25, error bars indicate the standard error of the mean; $^{*}p < 0.05$, $^{**}p < 0.01$, $^{****}p < 0.0001$). (**C**) The expression of $p27^{Kip1}$ is depicted as violin-bar conjugate plots, revealing increased expression with COL3A1 micropatterns under treatment (N = 3, n = 25, error bars indicate the standard error of the mean; $^{**}p < 0.01$, $^{***}p < 0.001$, $^{****}p < 0.0001$).

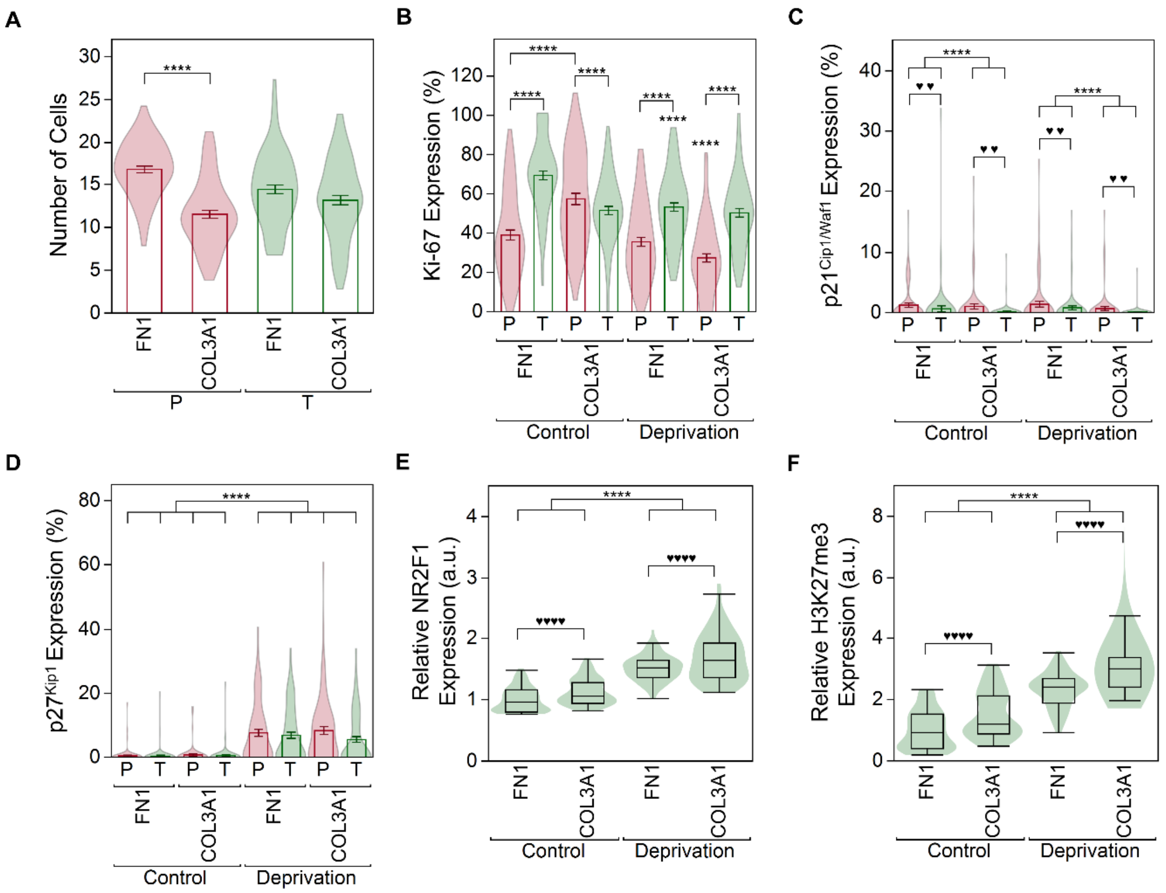


**Supplementary Figure 13: Effects of micropatterns and deprivation on SUM44 cells.**

(**A**) The attachment of SUM44-P and SUM44-T cells on FN1 and COL3A1 micropatterns is depicted as violin-bar conjugate plots, revealing differential attachment for the SUM44-P cells (N = 3, n = 25, error bars indicate the standard error of the mean; ****$p < 0.0001$). (**B**) The expression of Ki-67 is depicted as violin-bar conjugate plots (N = 3, n = 25, error bars indicate the standard error of the mean; ****$p < 0.0001$). (**C**) The expression of $p21^{Cip/Waf1}$ is depicted as violin-bar conjugate plots (N = 3, n = 25, error bars indicate the standard error of the mean; ****$p < 0.0001$, ♥♥$p < 0.05$ for the group). (**D**) The expression of $p27^{Kip1}$ is depicted as violin-bar conjugate plots, revealing increases for both SUM44-P and SUM44-T cells under deprivation conditions (N = 3, n = 25, error bars indicate the standard error of the mean; ****$p < 0.0001$). (**E**) The expression of NR2F1 in SUM44-T cells is depicted as violin-box conjugate plots (N = 3, n = 25, error bars indicate the standard error of the mean; ****$p < 0.0001$, ♥♥♥♥$p < 0.05$ for the group). (**F**) The expression of H3K27me3 in SUM44-T cells is depicted as violin-box conjugate plots (N = 3, n = 25, error bars indicate the standard error of the mean; ****$p < 0.0001$, ♥♥♥♥$p < 0.05$ for the group).

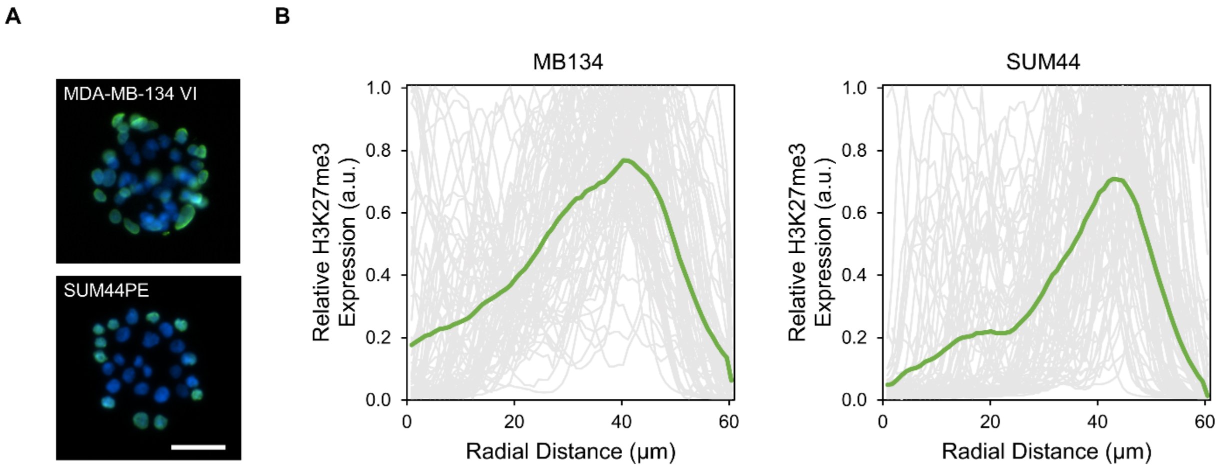


**Supplementary Figure 14: Radial expression of H3K27me3 in ILC cells.**

(**A**) Representative epifluorescence images of MB134-T and SUM44-T cells illustrate an outermost expression of H3K27me3 on FN1 micropatterns under control conditions. (**B**) The relative expression of the micropattern is plotted as a function of the radial distance emanating from the origin (the gray lines represent individual micropatterns and the green lines represent the mean; N = 3, n = 25). The scale bar is 50 µm.

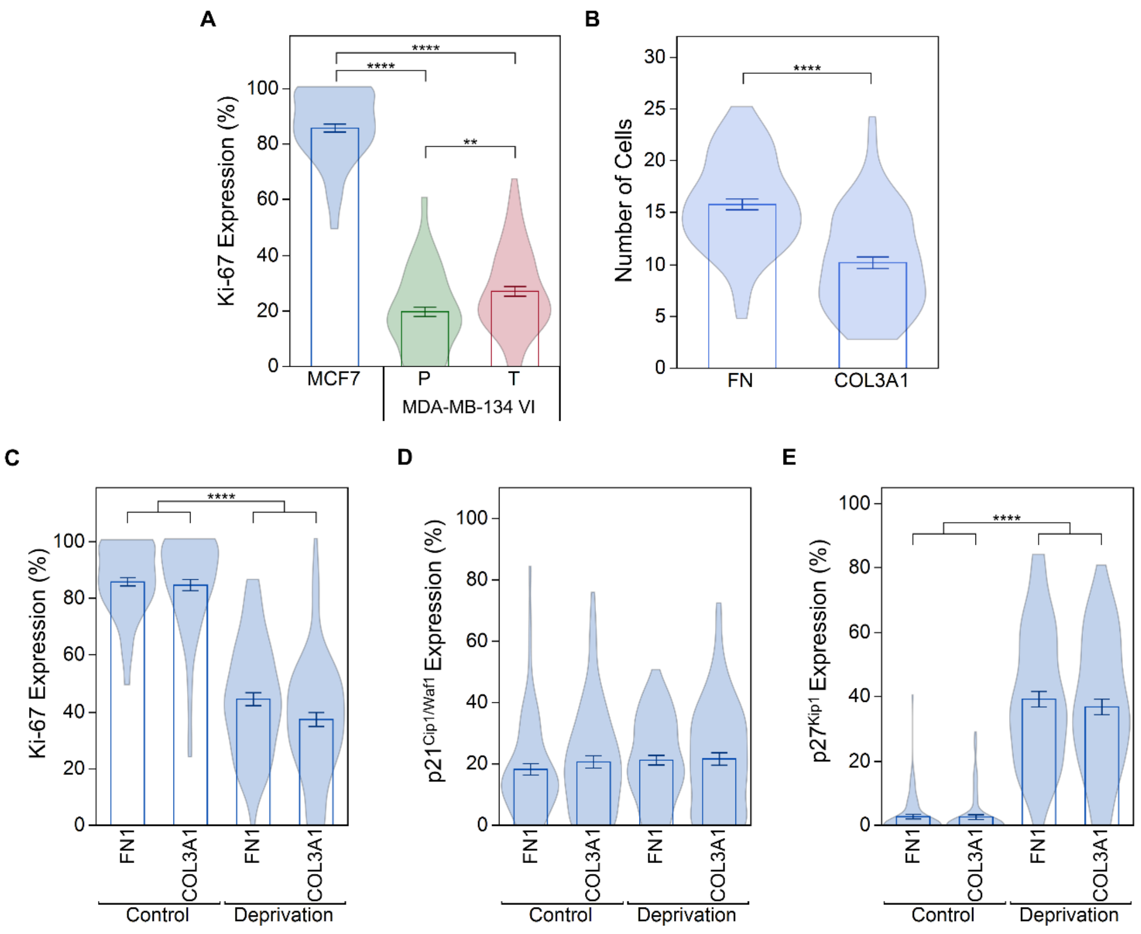


**Supplementary Figure 15: Effects of micropatterns and deprivation on MCF7 cells.**

(**A**) The expression of Ki-67 on the FN1 micropatterns under control conditions is compared across cell lines, with the highest expression observed in the MCF7 cells (N = 3, n = 25, error bars indicate the standard error of the mean; ***$p < 0.01$, ****$p < 0.0001$). (**B**) The attachment of MCF7 cells on FN1 and COL3A1 micropatterns is depicted as violin-bar conjugate plots, revealing an enhanced attachment to FN1 micropatterns (N = 3, n = 25, error bars indicate the standard error of the mean; ****$p < 0.0001$). (**C**) The expression of Ki-67 is depicted as violin-bar conjugate plots, illustrating a decrease under deprivation conditions (N = 3, n = 25, error bars indicate the standard error of the mean; ****$p < 0.0001$). (**D**) The expression of $p21^{Cip/Waf1}$ is depicted as violin-bar conjugate plots (N = 3, n = 25, error bars indicate the standard error of the mean). (**E**) The expression of $p27^{Kip1}$ is depicted as violin-bar conjugate plots, revealing increased expression under deprivation conditions (N = 3, n = 25, error bars indicate the standard error of the mean; $^{****}p < 0.0001$).

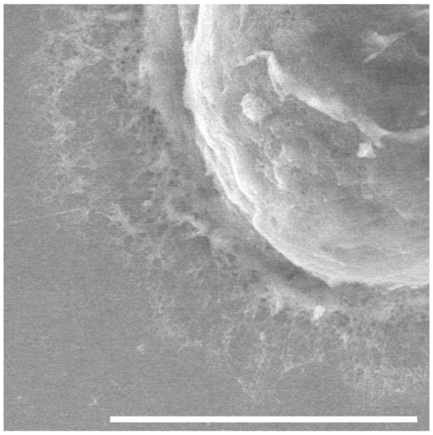

**Supplementary Figure 16: SEM image of MB134-P cell under deprivation conditions.**

A representative SEM image of an MB134-P cell on a COL3A1 micropattern under deprivation conditions shows an interaction with the patterned protein, albeit to a lesser extent than that of the MB134-T cells. The scale bar is 5 μm.

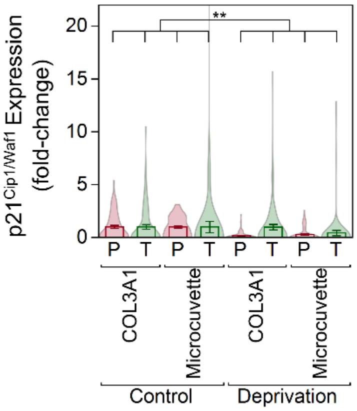


**Supplementary Figure 17: Comparing COL3A1 micropatterns and microcuvettes on p21$^{Cip1/Waf1}$ expression in MB134 cells.**

The expression of p21$^{Cip/Waf1}$ is depicted as violin-bar conjugate plots (N = 3, n = 25, error bars indicate the standard error of the mean; ***p* < 0.0001).

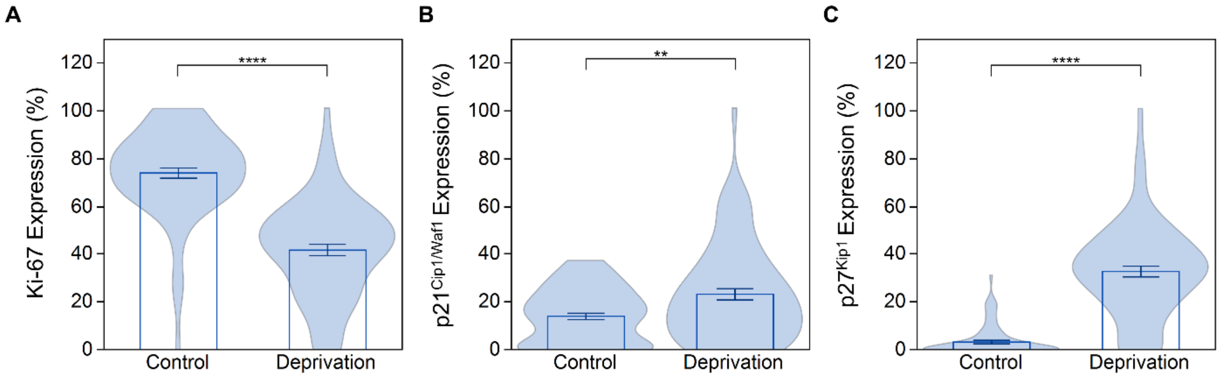


**Supplementary Figure 18: The effects of deprivation in MCF7 cells on the microcuvettes.** (**A**) The expression of Ki-67 is depicted as violin-bar conjugate plots, demonstrating a decrease under deprivation conditions (N = 3, n = 25, error bars indicate the standard error of the mean; ****$p$ < 0.0001). (**B**) The expression of $p21^{Cip/Waf1}$ is depicted as violin-bar conjugate plots, indicating an increase in expression under deprivation conditions (N = 3, n = 25, error bars indicate the standard error of the mean). (**C**) The expression of $p27^{Kip1}$ is depicted as violin-bar conjugate plots, revealing increased expression under deprivation conditions (N = 3, n = 25, error bars indicate the standard error of the mean; ****$p$ < 0.0001).
